# A one-stage approach for the spatio-temporal analysis of high-throughput phenotyping data

**DOI:** 10.1101/2023.01.31.526411

**Authors:** Diana M. Pérez-Valencia, María Xosé Rodríguez-Álvarez, Martin P. Boer, Fred A. van Eeuwijk

**Affiliations:** BCAM - Basque Center for Applied Mathematics, Mazarredo, 14 E48009 Bilbao, Basque Country, Spain; Departamento de Matemáticas, Universidad del País Vasco UPV/EHU, 48940 Leioa, Spain; CINBIO, Universidade de Vigo, Department of Statistics and Operations Research, 36310 Vigo, Spain; CITMAga, Centro de Investigación e Tecnoloxía Matemática de Galicia, 15782 Santiago de Compostela, Spain; Biometris, Wageningen University & Research, Wageningen, 6708 PB, The Netherlands

**Keywords:** Longitudinal analysis, mixed models, multidimensional P-splines, plant breeding, plant physiology, sparse structure

## Abstract

This work is motivated by the need to accurately estimate genetic effects over time when analysing data from high-throughput phenotyping (HTP) experiments. The HTP data we deal with here are characterised by phenotypic traits measured multiple times in the presence of spatial and temporal noise and a hierarchical organisation at three levels (populations, genotypes within populations, and plants within genotypes). We propose a feasible one-stage spatio-temporal P-spline-based hierarchical approach to model the evolution of the genetic signal over time on a given phenotype while accounting for spatio-temporal noise. We provide the user with appealing tools that take advantage of the sparse model matrices structure to reduce computational complexity. We illustrate the performance of our method using spatio-temporal simulated data and data from the PhenoArch greenhouse platform at INRAE Montpellier. In the plant breeding context, we show that information extracted for genomic selection purposes from our fitted genotypic curves is similar to those obtained using a comparable two-stage P-spline-based approach.

## 1 Introduction

Recent technological developments in plant breeding programmes have enabled data acquisition through high-throughput phenotyping (HTP) platforms. In this paper, we focus on HTP data in which a phenotypic trait (e.g., plant height, canopy cover, leaf area index, ear and tiller counts, canopy temperature, or indices related to water or chlorophyll content) is measured over time for a large number of plants and genotypes. In this setting, it is of primary interest to simultaneously model the longitudinal evolution of the genetic effect on a given phenotypic trait, while accounting for the temporal and spatial effects of environmental and design factors (van Eeuwijk et al., 2019). The challenge is to efficiently combine statistical and computational methods to deal with the size and complexity/dimensionality of the data while taking advantage of its spatio-temporal and hierarchical correlation structure.

In traditional agricultural experiments, a trait (generally yield) is measured only once, usually at the end of the experiment. For these experiments, it is well known that the phenotype of interest is spatially affected by micro-environmental factors, such as those that cause soil heterogeneity. To counterbalance spatial heterogeneity in field trials, various types of experimental designs are used (see, e.g., Brien et al., 2013; Hartung et al., 2019; Mead, 1997). In addition to using experimental design, spatial models can be used that allow to separate genetic and non-genetic effects properly (Besag & Higdon, 1999; Cullis & Gleeson, 1991; Durban et al., 2003; Gilmour et al., 1997; Green et al., 1985; Piepho & Williams, 2010; Rodríguez-Álvarez et al., 2018; Verbyla et al., 1999).

For recent HTP experiments, phenotypic traits are measured several times between, e.g., seed emergence and physiological maturity. Thus, the data provided by HTP experiments are spatio-temporal in nature. It means that their analysis requires modelling strategies that, in addition to the spatial component, account for temporal dynamics. Research in this area is increasing, but the data complexity asks for novel methods. One possibility is to use stagewise approaches (Kar et al., 2020; Pérez-Valencia et al., 2022; Roth et al., 2021; van Eeuwijk et al., 2019). Stage-wise proposals have the advantage of being computationally manageable, but the problem relies on the loss of information between and within stages. For instance, for the two-stage P-splines-based approach proposed by Pérez-Valencia et al. (2022), spatial heterogeneity is not shared across time when correcting for environmental factors in the first stage, and uncertainty is lost between stages (weights are used to propagate error from the first to the second stage). Kar et al. (2020) and van Eeuwijk et al. (2019) first estimate spatially adjusted genotypic means per time point and, in a second stage, they model the genotypic signal over time independently for each genotype ignoring the hierarchical data structure that is shared between genotypes within populations and between plants within genotypes.

It is therefore of interest to develop approaches that allow modelling the spatial and temporal genetic and non-genetic variation in one stage. In this setting, Verbyla et al. (2021) have proposed modeling the genetic effects over time using factor analytic models and use smoothing splines to model the non-genetic/residual effects over time and space. Nevertheless, the authors report that their work is a “proof of concept” in the sense that fitting the models is very time-consuming. In fact, they ask for scalable software. To overcome these computational issues and take advantage of all the available information given by the data structure, we propose a one-stage spatio-temporal P-spline hierarchical curve data model. In particular, we generalise the two-stage modelling strategy presented in Pérez-Valencia et al. (2022) to a full and one-stage spatio-temporal approach. We use the SpATS (Spatial Analysis of field Trials with Splines, Rodríguez-Álvarez et al., 2018) model as the base model and extend it to the spatio-temporal case, considering a three-level hierarchical data structure (populations, genotypes within populations, and plants within genotypes). Additionally, we exploit the connection between P-splines and linear mixed models and the sparse structure of the matrices involved in the model to improve computational time. As results, we obtain estimated curves and their derivatives at the three levels of the hierarchy (populations, genotypes and plants). We use these curves to extract different time-independent characteristics (intermediate traits) that can be used in further analysis (see, e.g., van Eeuwijk et al., 2019 and Moreira et al., 2020).

The rest of the paper is organised as follows: In Sect. 2, we present our proposal and describe the estimation methods and some computational aspects. We also assess the performance of our approach through simulated data and compare its results with those obtained using a two-stage approach (Pérez-Valencia et al., 2022). In Sect. 3, we again compare the two proposals by analysing real HTP data from the INRAE greenhouse platform in Montpellier. Finally, in Sect. 4, we discuss the advantages and limitations of our approach.

## 2 One-stage approach

The HTP data we consider in this paper consist of a phenotypic trait measured several times at individual experimental units (hereafter referred, for simplicity, as plants). We assume that the genotypes are allocated to the experimental units following an experimental design, that no other treatments are assigned, and that the experimental units can be mapped to a coordinate system defined in terms of *R* rows and *C* columns. Furthermore, we allow for population structure (i.e., families or panels of genotypes). Thus, the data consist of a sample of plant curves (time-series) that can be grouped by two (nested) factors (genotypes and populations). Following the notation in Brumback and Rice (1998) and Xu et al. (2021), we denote as *y_i_*(*t*) the observed phenotypic trait for the *i*th plant (*i* = 1,…, *M*) at time *t* (*t* ∈ {*t*_1_,…, *t_n_*}). We use *p*, *g*, *r* and *c* for indices of population, genotype, and row and column position, respectively (*p* = 1,…, *K*; *g* = 1,…, *L*; *r* = 1,…, *R*; *c* = 1,…, *C*). With a slight abuse of notation, we use *p*(*i*), *g*(*i*), *r*(*i*) and *c*(*i*) to denote the population, genotype, and row and column position of the ith plant, respectively. Finally, we define *l_p_* = #{*g* | *p*(*g*) = *p*} as the number of genoypes with population *p*, and *m_g_* = #{*i* | *g*(*i*) = *g*} as the number of plants with genotype *g* (i.e., 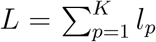 and 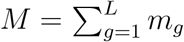). Note that we can have a different number of genotypes per population and/or a different number of replicates (plants) per genotype. Besides, we assume that all plants are measured at the same *n* time points ({*t*_1_,…, *t_n_*}), yet missing observations for a plant curve are also allowed.

### 2.1 Spatio-temporal P-spline hierarchical curve data model

Our approach builds upon the spatial SpATS model proposed by Rodríguez-Álvarez et al. (2018). In particular, if *y_i_* is the phenotypic trait for the ith plant for a specific time point (for simplicity, we omit here the dependence on time), the SpATS model we consider is as follows

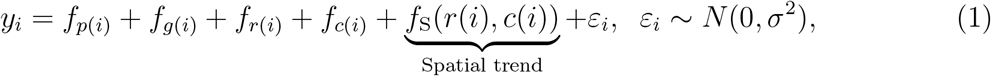

where *f_p_* is the fixed effect coefficient for population *p*, *f_g_* is the random effect coefficient for genotype 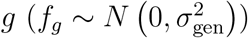, and *f_r_* and *f_c_* are random effect coefficients for row *r* and column *c*, respectively 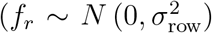 and 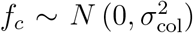; they are included to account for design factors). Finally, *f_s_*(*r,c*) (where S stands for Spatial) is a two-dimensional smooth function, defined over the row and column positions, that simultaneously accounts for the spatial (local and global) trend variation across both directions. This smooth function is constructed with tensor-product P-splines (Eilers & Marx, 1996, 2003). As discussed in Rodríguez-Álvarez et al. (2018), this function can be decomposed in the following nested ANOVA-way

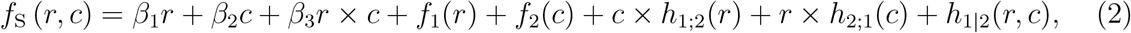

where *f*_1_(*r*) and *f*_2_(*c*) are smooth (non-linear) trends (main effects) along the rows and columns, respectively; *c* × *h*_1;2_(*r*) and *r* × *h*_2;1_(*c*) are linear-by-smooth interaction trends (varying coefficient surface terms; for instance *c* × *h*_1;2_(*r*) are linear trends in the columns (*c*) but with slopes (*h*_1;2_(*r*)) that change smoothly along the rows); and *h*_1|2_(*r, c*) is a pure smooth-by-smooth interaction trend jointly defined over the row and column directions. (for a detailed discussion, see Rodríguez-Álvarez et al., 2018).

By taking the SpATS model (1) as the base model, we now extend it to the spatiotemporal case by allowing all effects to vary with time, i.e.,

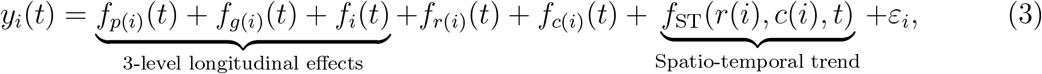

where *ε_i_*(*t*) has been decomposed such that *ε_i_*(*t*) = *f_i_*(*t*) + *ε_i_*, that is, we capture the (structured) temporal trend of each plant in *f_i_*(*t*), while *ε_i_* is pure random noise, i.e., *ε_i_* ~ *N*(0, *σ*^2^). The interpretation of each component in model (3) is as follows. *f_p_*(*t*) is the time-varying effect coefficient for population *p*, *f_g_*(*t*) is the time-varying random effect coefficient for genotype *g* (it measures deviations from the population effect to which the genotype belongs to), *f_i_*(*t*) is the time-varying random effect coefficient for plant *i* (it measures deviations from the genotype effect to which the plant belongs to), and *f_r_*(*t*) and *f_c_*(*t*) are time-varying random effect coefficients for row *r* and column *c*, respectively. Finally, *f*_ST_(*r*, *c*, *t*) is a spatio-temporal (ST) three-dimensional surface defined over rows, columns and time. This three-dimensional surface accounts for spatial trend variations, but it allows these spatial trends to change with time.

To model (and estimate) the time-varying effects in model (3), in this paper we assume that all these effects vary smoothly along time. Furthermore, we propose using P-splines for that purpose (Eilers & Marx, 1996, 2003), i.e., we combine B-spline basis functions on equidistant knots and a discrete difference penalty on the (regression) coefficients to ensure smoothness. In the next section, we present in detail the specification of model (3) and discuss its estimation as well as some computational aspects.

### 2.2 Model specification, estimation and computational aspects

Each univariate function in (3), *f_p_*(*t*), *f_g_*(*t*), *f_i_*(*t*), *f_r_*(*t*), and *f_c_*(*t*), is modelled as a linear combination of cubic B-spline basis functions, and *f*_ST_(*r*, *c*, *t*) using the tensor product of three marginal cubic B-spline bases. As said before, in P-splines smoothness is ensured by imposing a (second-order) difference penalty on the regression coefficients. The influence of the penalty (i.e., the amount of smoothness) is determined by one smoothing parameter in the univariate case, and by three smoothing parameters in the three-dimensional case, i.e., we consider anisotropy (details are given in Web Appendix A, but we refer the reader to Eilers & Marx, 2021; Rodríguez-Álvarez et al., 2015, and references therein for a more in-depth presentation).

For estimation, we use the connection between P-splines and linear mixed models (Currie & Durban, 2002; Currie et al., 2006; Lee & Durban, 2011; Wand, 2003). Here, the smooth functions are decomposed (reparameterised) in two components: one whose coefficients are not penalised (in the mixed model framework, these coefficients are considered as fixed), and one whose coefficients are penalised (and thus considered as random). Besides, smoothing parameters are “replaced” by variance parameters, and estimated using restricted maximum likelihood (REML; Patterson and Thompson, 1971). In our setting, however, for *f_g_*(·), *f_i_*(·), *f_r_*(·) and *f_c_*(·) the unpenalised coefficients (intercept and slope; we use second-order difference penalties) are also assumed to be random. There are two reasons for this decision. On the one hand, we treat genotypes and plants as random samples. On the other hand, it avoids identifiability problems that arise with fixed effects for nested ANOVA models (Brumback & Rice, 1998). In summary, the modeling strategy we follow implies that there is one variance parameter per population, and three variance parameters (associated, respectively, with the intercept, the slope and the non-linear/penalised/smooth effect) for genotypes, plants, rows and columns. The more technical details are given in Web Appendix A, but under this framework, each univariate function in model (3) is expressed as follows (for one plant)

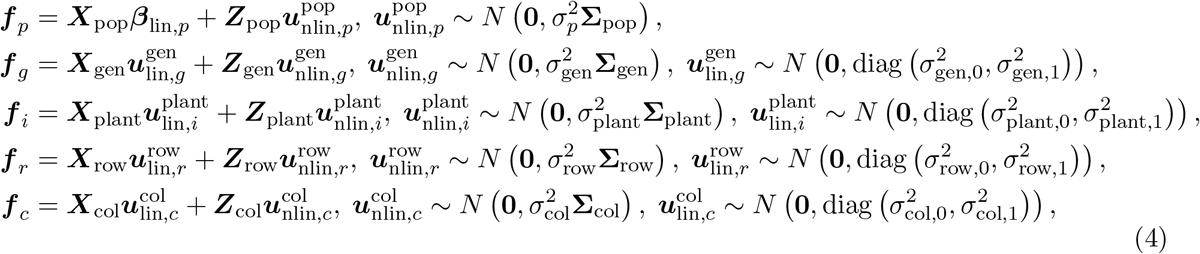

where ***f**_p_* = (*f_p_*(*t*_1_),…, *f_p_*(*t_n_*))^*T*^, ***β***_lin,*p*_ = (*β*_lin,*p*,0_, *β*_lin,*p*,1_)^*T*^, ***X***_pop_ = [**1**_*n*_ | ***t***], **1**_*n*_ is a column vector of ones of length *n*, ***t*** = (*t*_1_,…, *t_n_*)^*T*^ (to be more precise, ***X***_pop_ is obtained as described in Wood et al., 2013, see also Web Appendix A), 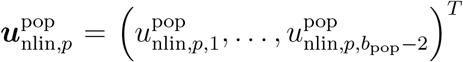 and (***Z***_pop_)_*lj*_ = *z*_pop,*j*_(*t*_l_) with {*z*_pop,*j*_(·): 1 ≤ *j* ≤ *b*_pop_ − 2} being a set of basis functions. These basis functions, as well as the variance-covariance matrix of the random vector 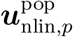, **Σ**_pop_, are obtained from the connection between P-splines and linear mixed models (see Web Appendix A for technicalities). All of the above also applies to the other univariate functions in (4), but to avoid notational overhead we omit the details.

With respect to the spatio-temporal three-dimensional surface in model (3), *f*_ST_(*r,c,t*), we follow the ideas in Lee and Durban (2011) and Rodríguez-Álvarez et al. (2015) for its mixed model reparameterisation (for details, see also Web Appendix A). Before proceeding, it is worth indicating that *f*_ST_(*r*, *c*, *t*) can also be decomposed in a nested ANOVA-way similar to the one shown for the two-dimensional case (see (2))

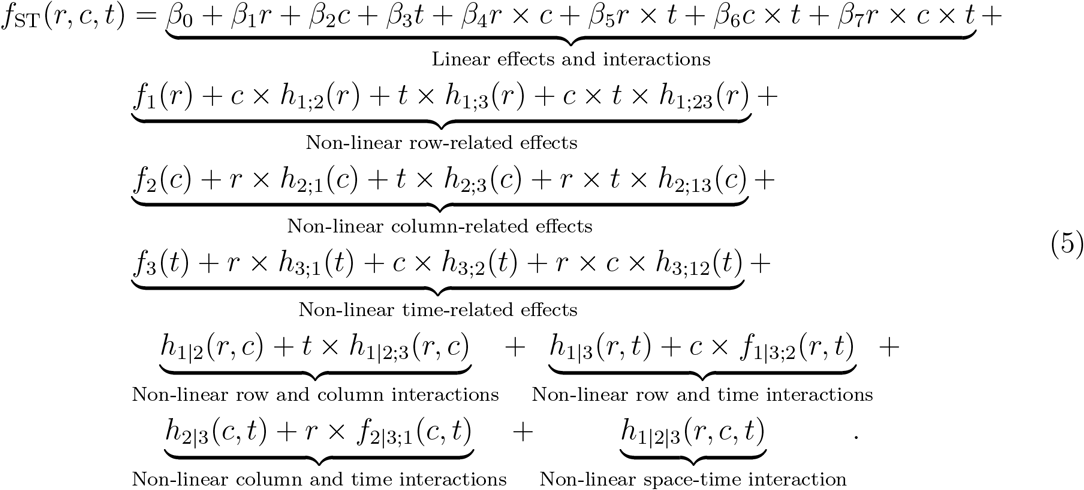

In contrast to the two-dimensional case, more components are now present (there are three variables involved: rows, columns and time), yet their interpretation is similar. For instance, *c* × *t* × *h*_1;23_(*r*) are linear interaction trends in the columns (*c*) and time (*t*), but with slopes (*h*_1;23_(*r*)) that change smoothly along the rows. Although the decomposition shown in (5) won’t be further exploited in the application, it will be helpful when interpreting the (mixed model) design matrices associated with the three-dimensional surface. In addition, it reveals that there are several components which are confounded with the population effect, namely, the intercept *β*_0_, the linear effect along time *β*_3_*t* and the non-linear (smooth) main effect along time *f*_3_(*t*). These are removed from the specification that follows. In a similar fashion to the univariate case, each non-linear (smooth) function in (5) (i.e, *f*_{·}_ and *h*_{·}_) is specified as a linear combination of (tensor-product) basis functions. In matrix notation, the mixed model form of the spatio-temporal three-dimensional surface is

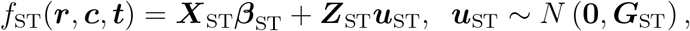

where *f*_ST_(***r***, ***c***, ***t***) = (*f*_ST_(*r*(1), *c*(1), *t*_1_), *f*_ST_(*r*(1), *c*(1), *t*^2^),…,*f*_ST_ (*r*(*M*), *c*(*M*), *t_n_*))^*T*^, and

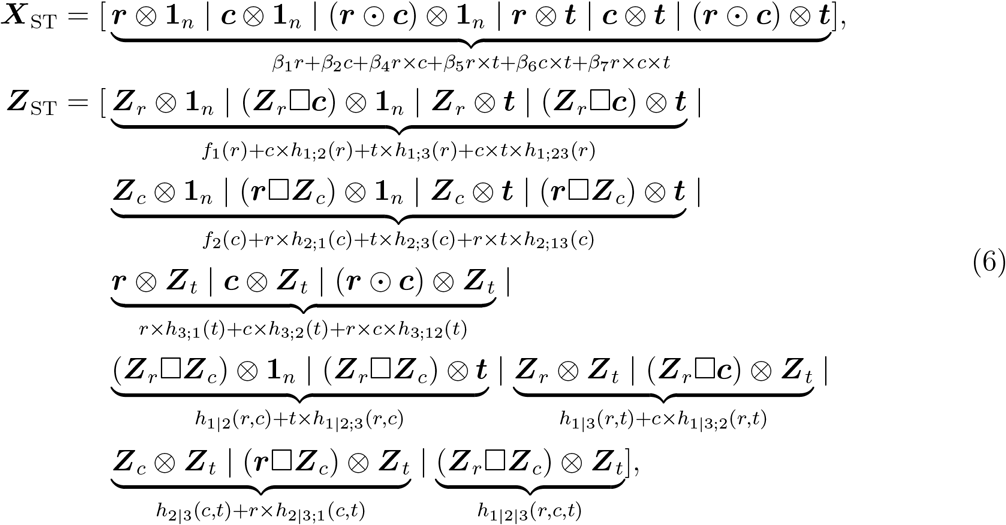

where ***r*** = (*r*(1), *r*(2),…, *r*(*M*))^*T*^, ***c*** = (*c*(1), *c*(2),…, *c*(*M*))^*T*^, ⊗ denotes the Kronecker product, ⊙ the element-wise (Hadamard) product, and □ the “box” product (the face-splitting product or row-wise Kronecker product, Eilers et al., 2006; Slyusar, 1999), i.e., 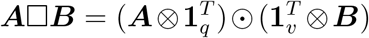, where ***A*** and ***B*** are two (generic) matrices of dimension *m* × *q* and *m* × *v*, respectively. Finally, (***Z**_r_*)_*ij*_ = *z_r,j_*(*r*(*i*)), (***Z**_c_*)_*ij*_ = *z_c,j_*(*c*(*i*)) and (***Z**_t_*)_*lj*_ = *z_t,j_*(*t_l_*), where, as for the univariate case, {*z_r,j_*(·): 1 ≤ *j* ≤ *b_r_* − 2}, {*z_c,j_*(·): 1 ≤ *j* ≤ *b_c_* − 2}, and {*z_t,j_*(·): 1 ≤ *j* ≤ *b_t_* − 2} are sets of basis functions obtained from the connection between P-splines (with second-order difference penalties) and linear mixed models. Regarding the vectors of fixed and random effect coefficients, we have

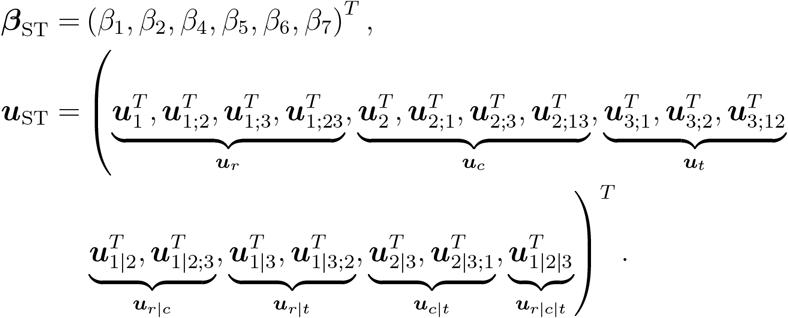

Note that for the vector of random effects, ***u***_ST_, we have a total of 18 sets of random effects, each associated with one block in ***Z***_ST_ (i.e, with one non-linear/smooth function in (5)). These sets can be further grouped into 7 larger sets. The first three, ***u**_r_*, ***u**_c_* and ***u**_t_*, correspond to univariate non-linear/smooth effects along the rows, columns and time, respectively, and ***u***_*r*|*c*_, ***u***_*r*|*t*_ and ***u***_*c*|*t*_ to bivariate non-linear/smooth interactions between rows and columns, rows and time, and columns and time, respectively. Finally, ***u***_*r*|*c*|*t*_ correspond to the trivariate non-linear/smooth interaction between rows, columns and time. With these 7 sets in mind, the precision matrix (i.e., the inverse of variance-covariance matrix) associated with ***u***_ST_, is a block diagonal matrix, with 7 blocks, each related to each set of random effects

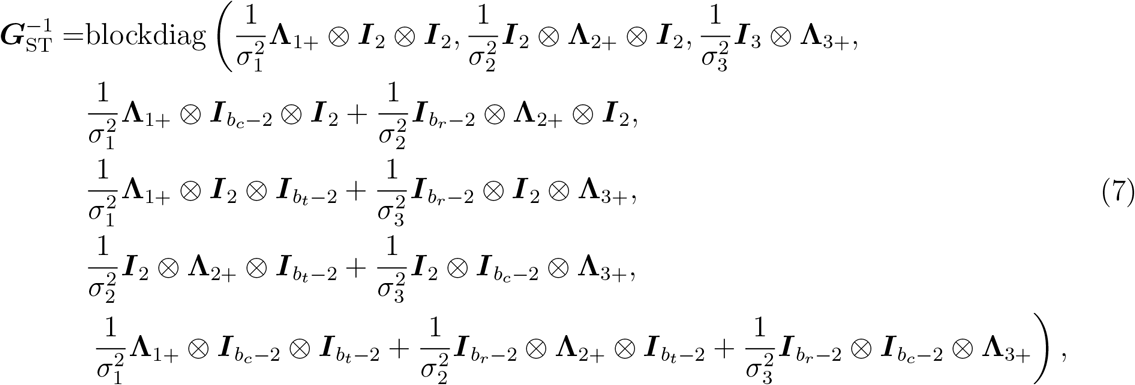

where the specific form of **Λ**_*j*+_ (*j* = 1, 2, 3) is given in Web Appendix A (once again it is obtained from the connection between P-splines and linear mixed models) and ***I**_d_* denotes the identity matrix of dimension *d*. As can be observed, 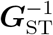 depends on three variance parameters, 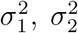, and 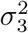 they are responsible for controlling the smoothness along the rows, columns and time, respectively.

With all ingredients introduced before (i.e., the specifications in equations (4) and (6)), in matrix form model (3) is expressed, for all *M* plants, as

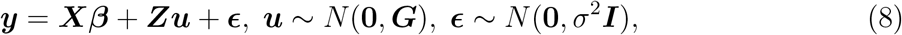

where

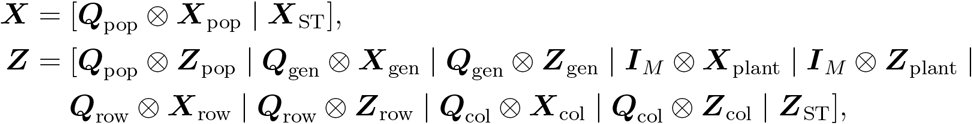

where 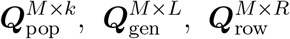 and 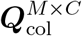 are matrices assigning, respectively, plants to populations, plants to genotypes, plants to row locations, and plants to column locations (the super-indices indicate their dimension). Furthermore, 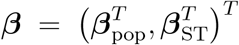 and 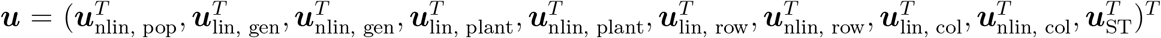, where

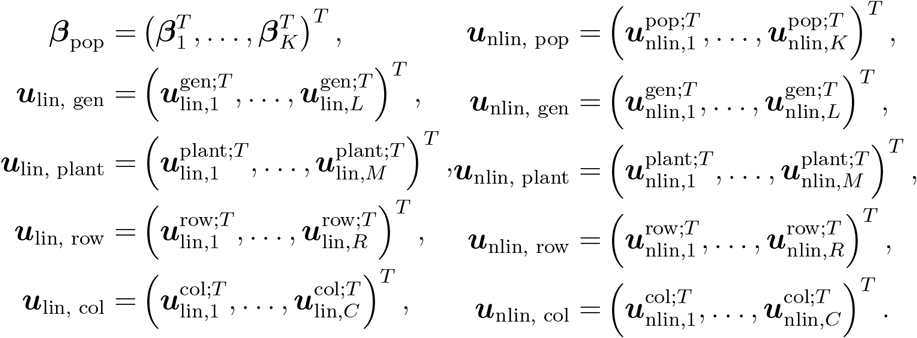

Finally, the variance-covariance matrix ***G*** is a block diagonal matrix given by

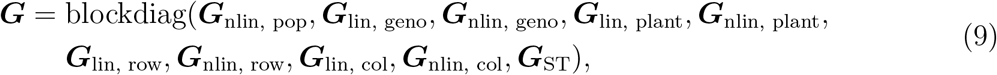

where

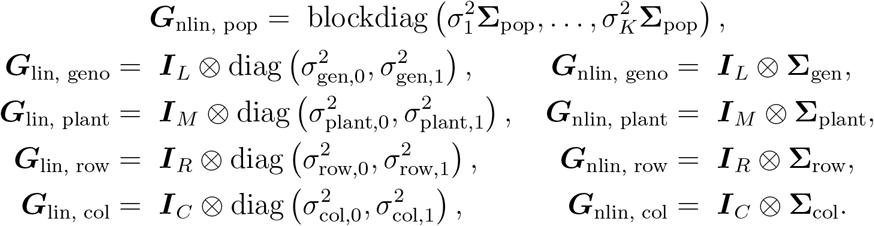

We note that the variance-covariance (9) has a non-standard form, with the last block in ***G***_ST_ depending on three variance parameters in a non linear way. More precisely, in our case what is linear in the (inverse of the) variance parameters is the precision matrix, ***G***^−1^ and not the variance-covariance matrix, ***G***. This feature precludes estimation of model (8) using standard mixed models software, and specialised methods are required (e.g., Rodríguez-Álvarez et al., 2019; Rodríguez-Álvarez et al., 2015). Besides, the data size produced by HTP experiments, together with the large number of regression coefficients associated with P-spline-based methods, make the problem computationally challenging. To keep the computational effort manageable, we implemented our own code in the `R` language (R Core Team, 2021) (provided along with the paper), and use the so-called SOP (Separation of Overlapping Precision matrices) method proposed by Rodríguez-Álvarez et al. (2019). Empirical best linear unbiased estimates (BLUE) and predictors (BLUP) are obtained by the solution of Henderson’s mixed model equations (Henderson, 1963), and variance parameters are estimated using REML. Here, we use the `R`-package `LMMsolver` (Boer & van Rossum, 2022) to calculate partial derivatives of the REML log-likelihood in an efficient way. The use of this package allows us to further reduce the computational burden by exploiting the sparse structure of the matrices involved in the model by using the automated differentiation of Cholesky algorithm proposed by Smith (1995). Finally, computational calculations are further sped up by taking advantage of the array structure of the data through Generalised Linear Array Models (GLAM; Currie et al., 2006).

### 2.3 Model performance assessment through simulation

In this section, we evaluate the performance of the proposed one-stage model and compare the results with those obtained with the two-stage approach described in Pérez-Valencia et al. (2022). When applying these two approaches to real data sets, two problems arise: (i) the phenotype of interest is not measured at the population and genotype levels but only at the plant level; it makes evaluating models’ performance at these levels difficult, and (ii) the approaches seem to be sensitive to the dimension of the B-spline bases used at each level of the hierarchy. We present a data generating model and a simulation experiment to study the above-mentioned problems.

#### 2.3.1 Data generating model

Following the same notation introduced in the previous section, we simulate HTP data assuming the following three-level nested hierarchical model

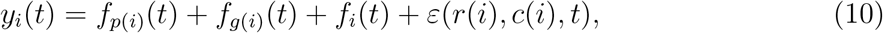

where 1 ≤ *g* ≤ *L*, 1 ≤ *i* ≤ *M*, 1 ≤ *r* ≤ *R* and 1 ≤ *c* ≤ *C*. For simplicity, here we consider only one population (*K* = 1), and *M* = *R* × *C* plants (i.e., the number of plants corresponds to the number of spatial locations, with a 1:1 correspondence). In what follows we assume that the plants are located on the R × C grid such that the first plant (*i* = 1) is in the first row and column position (i.e., *r*(1) = 1 and *c*(1) = 1), and the last plant (*i* = *M*) is in the last row and column position of the grid (i.e., *r*(*M*) = *R* and *c*(*M*) = C), and plants are ordered by rows (see Web Figure 1). We note that the data generating model, equation (10), is independent from the statistical model, equation (3), used for the analysis. The difference between the two models lies in how the spatio-temporal (stochastic) component is incorporated. In model (10), we have spatio-temporal correlated noise, *ε*(*r*, *c*, *t*), while in model (3), we have a spatio-temporal three-dimensional surface, *f*_st_(*r*, *c*, *t*), and the pure random noise, *ε_i_*. Besides, model (10) does not consider the row and column random effects. In particular, the data are generated from the population to the plant level in five steps. For brevity, these steps are described in detail in Web Appendix B.1, but we provide Figure 1 to graphically summarise the procedure and the kind of curves that are obtained at each step.

**Figure 1:**
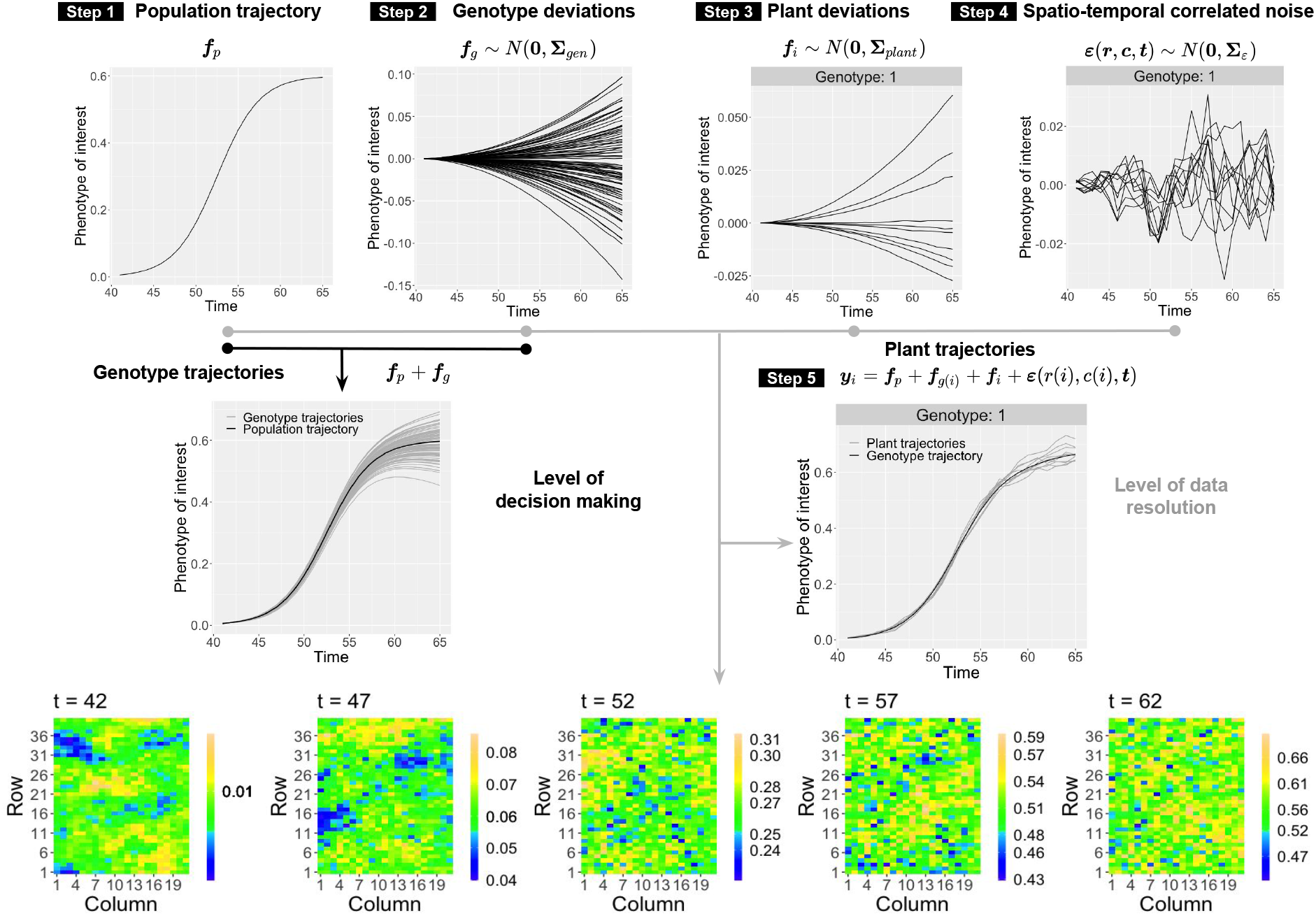
For the simulation study: Data generating strategy.

#### 2.3.2 Simulation scenarios and set-up

Data is simulated under eight different scenarios given by the combination of the levels of three factors, each with two levels: (i) the between genotype (deviation) variability, 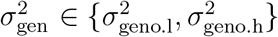; (ii) the between plant (deviation) variability, 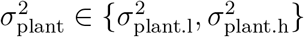; and, (iii) the number of replicates per genotype, *m_g_* ∈ {3,10}. For 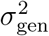 and 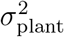, l stands for low and h for high, that is, 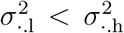. Each scenario is denoted as (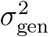, 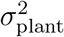, *m_g_*), and they represent different levels of heritability (see, e.g., Rodríguez-Álvarez et al., 2018, for a definition of heritability). For instance, scenarios with *m_g_* = 10 replicates per genotype have higher heritability than those with *m_g_* = 3 replicates; and the scenario with the highest heritability is (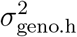, 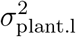, 10), that is, the scenario where the between genotype variability is higher than the between plant variability (see Web Figure 2 for a comparison of the heritability among simulation scenarios). For each scenario, 100 datasets are generated and the simulation settings are described in Web Table 1.

**Figure 2:**
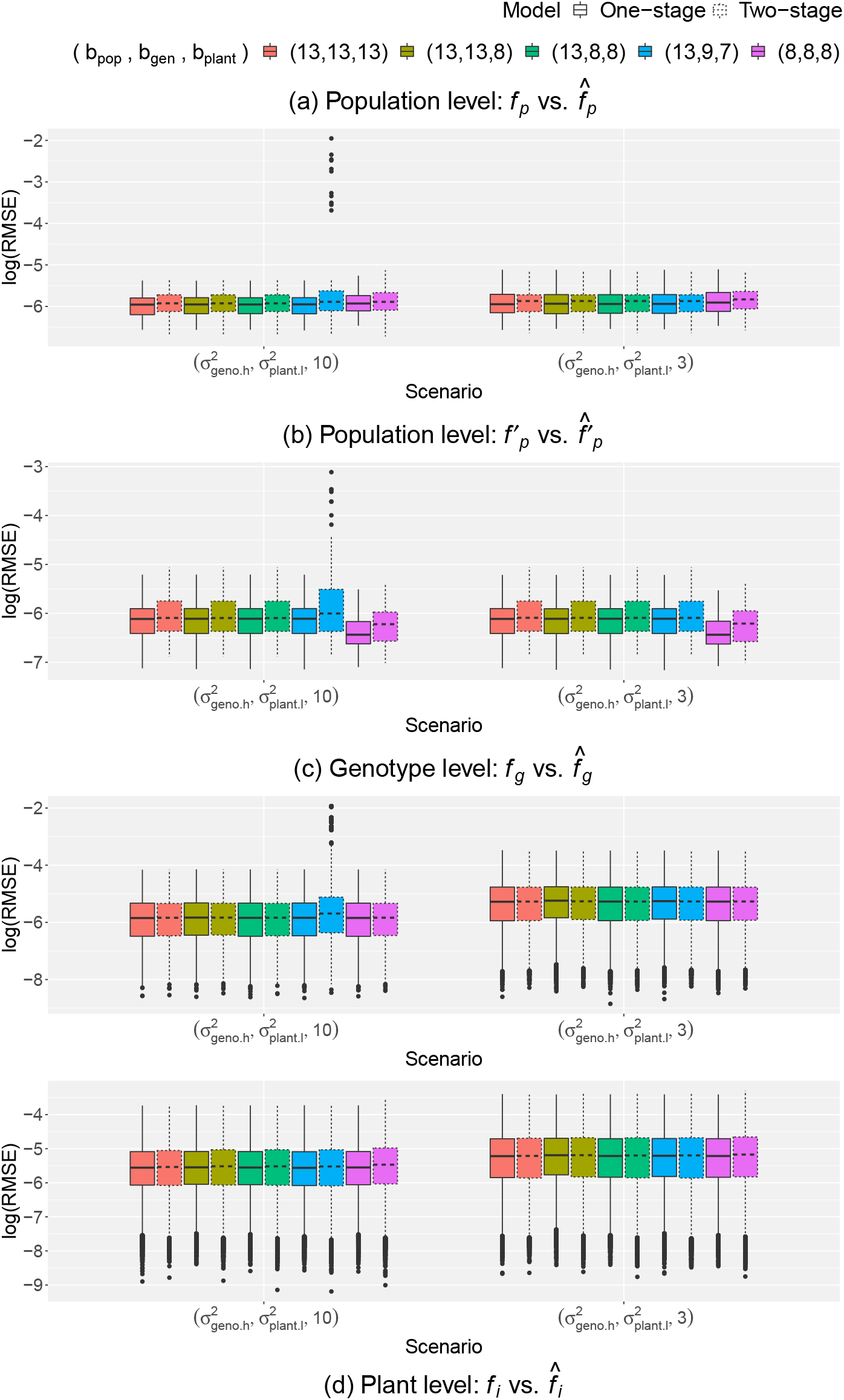
Simulation results: comparison of the simulated and estimated (a) population trajectories, (b) first-order derivative of the population trajectories, (c) genotype deviations curves, and (d) plant deviation curves for two of the eight scenarios of data simulation (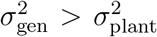 and *m_g_* = 3, 10), using the one- and two-stage approaches, and five B-spline basis configurations (*b*_pop_, *b*_gen_, *b*_plant_) at population, genotype and plant level, respectively.

For the one-stage approach, we fit model (3) with *b*_row_ = *b*_col_ = 13 B-splines for the random row, *f_r_*, and column, *f_c_*, effects (see (4)), and *b_r_* = *b_c_* = *b_t_* = 13 for the spatio-temporal smooth function, *f*_ST_ (see (5)). In addition, we compare the performance of five different configurations for the dimensions of the B-spline bases for the hierarchical components, *f_p_*, *f_g_* and *f_i_* (see (4)). In particular, we consider (*b*_pop_, *b*_gen_, *b*_plant_) ∈ {(13, 13, 13), (13, 13, 8), (13, 8, 8), (13, 9, 7), (8, 8, 8)}. The first and fifth configuration aim to assess model’s flexibility and over-fitting, the second and third configuration evaluate the possible impact of considering different bases dimensions at different levels of the hierarchy, and the fourth configuration assesses model performance under non-nested bases. In this context, nested bases refers to B-spline bases such that the space spanned by ***B***_plant_ and ***B***_gen_ are subsets of the space spanned by ***B***_pop_ (for an example see Web Figure 3, and for more technical details we refer to Lee et al., 2013). To make the one and two-stage approaches comparable, we set the dimensions of the B-spline bases of the model components that are common to the two approaches to the same values. Consequently, for the first stage of the two-stage approach, we fit a SpATS model separately for each time point, with *b_r_* = *b_c_* = 13 B-splines for the two-dimensional smooth function, fS (see also (1) and (2)). For the second stage, we fit the P-spline hierarchical curve data model using the five different configurations previously described for the one-stage approach (for details on the two-stage approach, we refer the reader to Pérez-Valencia et al., 2022). We used the logarithm of the root mean square error (log(RMSE)) as performance measure to compare the simulated and the estimated curves.

**Figure 3:**
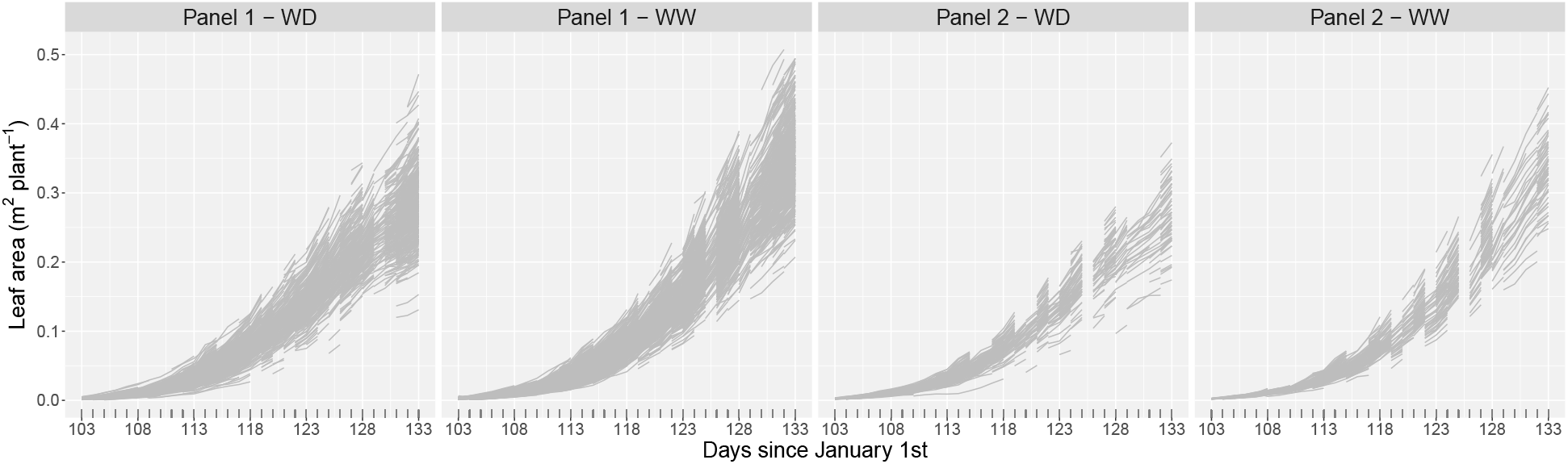
Evolution over time of the raw leaf area by population for data from the PhenoArch platform.

#### 2.3.3 Simulation results

We obtained estimated curves at the three levels of the hierarchy: (1) population trajectories 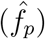 and respective first-order derivatives 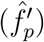, (2) genotype-specific deviations 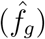 and respective trajectories 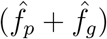, and (3) plant-specific deviations 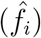 and respective trajectories 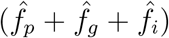. We focus here on the results for the population trajectories and their first-order derivatives as well as for the genotype- and plant-specific deviations. For brevity, we present results for two of the eight scenarios that correspond to the highest heritability, i.e., (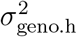, 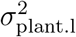, 10) and (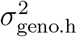, 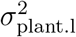, 3) (Web Figures 4, 5 and 6 depict the results for the remaining scenarios). Figure 2 shows the log(RMSE) for both scenarios, approaches (one-stage in dotted border boxplots, and two-stage in continuous border boxplots), and the five different B-splines basis configurations (in colours). Results at population and genotype levels, Figures 2(a), 2(b) and 2(c), show that both approaches perform similarly except for the non-nested basis (i.e., *b*_pop_ = 13, *b*_gen_ = 9, *b*_plant_ = 7; results in blue) and *m_g_* = 10 replicates, where the one-stage approach performs better. Results for the population trajectories, Figure 2(a), and plant deviations, Figure 2(d), show that, for the two-stage approach, the lesser the flexibility (i.e., *b*_pop_ = 8, *b*_gen_ = 8, *b*_plant_ = 8; results in purple), the worse the performance. However, this configuration is the one performing the best for the first-order derivative of the population trajectories, Figure 2(b) (results in purple). The largest differences between the one- and two-stage approaches are observed at the population level, with the one-stage approach performing better. For the population trajectories, Figure 2(a), differences between both approaches are more evident for scenarios with *m_g_* = 10 replicates. For the first-order derivative of the population trajectory, Figure 2(b), differences remain for all scenarios and all B-spline basis configurations. In general, the more replicates the better the performance of the two approaches, but the number of replicates affects more (the difference is bigger) at the genotype level. All in all, for most studied conditions slight differences appear between the two approaches. For the scenarios with the largest differences, the one-stage approach consistently outperformed the two-stage approach.

**Figure 4:**
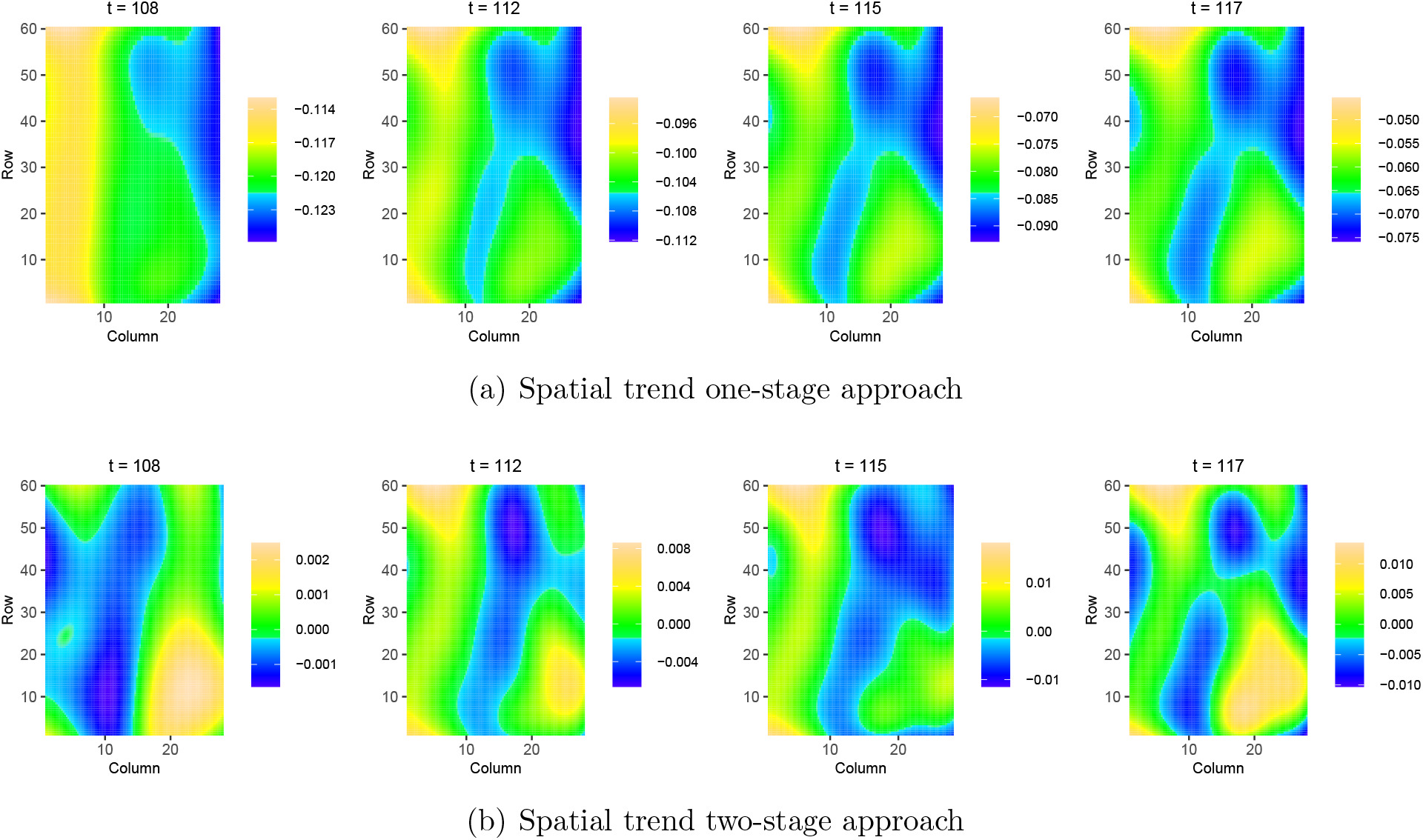
Results for the PhenoArch platform: spatial trend of the (a) one-stage approach, and (b) two-stage approach, at four different measurements times (*t* = 108, 112, 115, 117 DOY). The colour scale is different for each time point.

**Figure 5:**
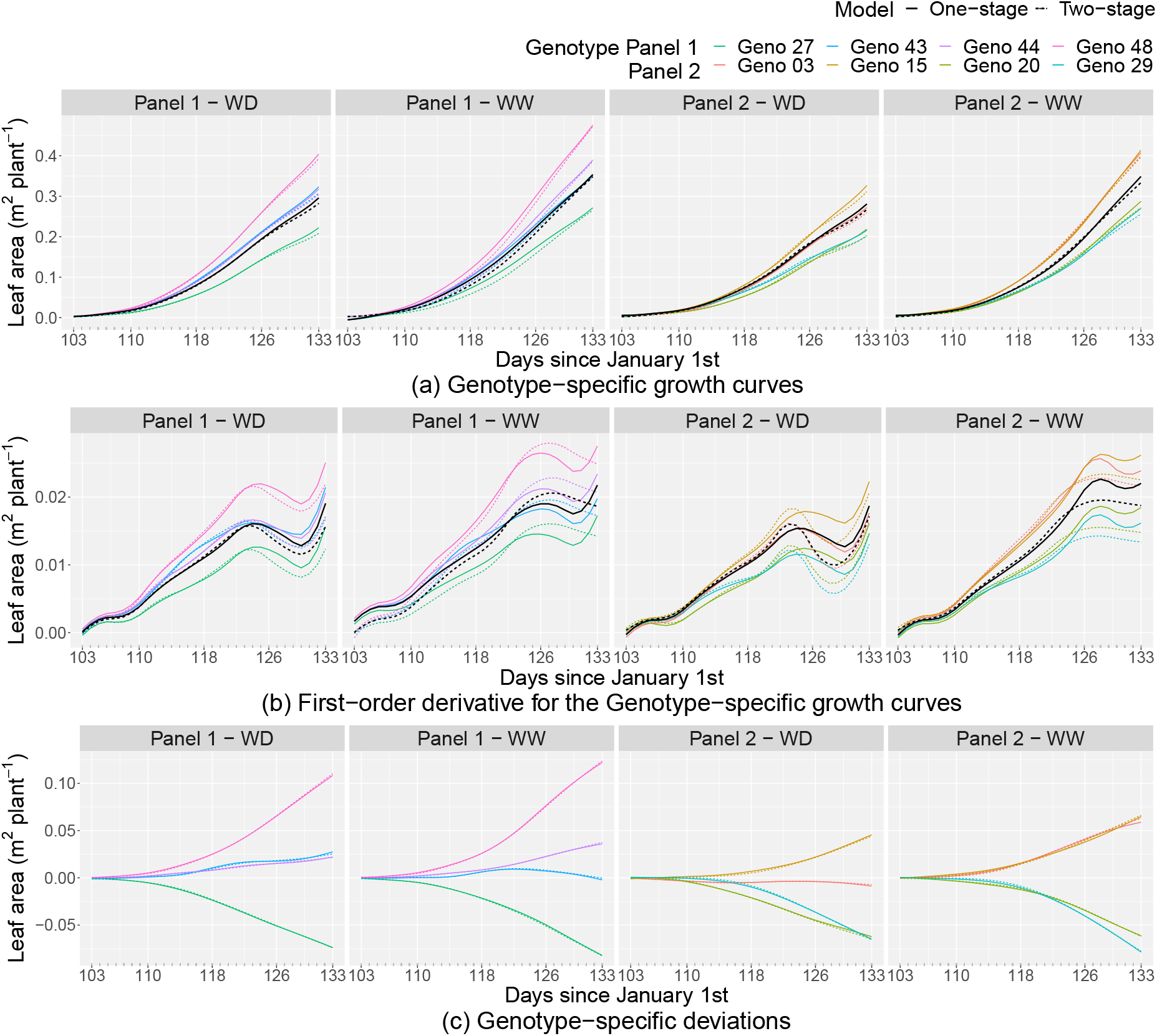
Results for the PhenoArch platform at the genotype level (for four genotypes by panel, as illustration) for the one-stage (continuous lines) and two-stage (dotted lines) approaches, separately for each panel-by-water regime combination: (a) estimated genotypespecific growth curves, (b) estimated first-order derivative for the genotype-specific growth curves, and (c) estimated genotype-specific deviations. In Figures (a) and (b) black lines represent curves at population level.

**Figure 6:**
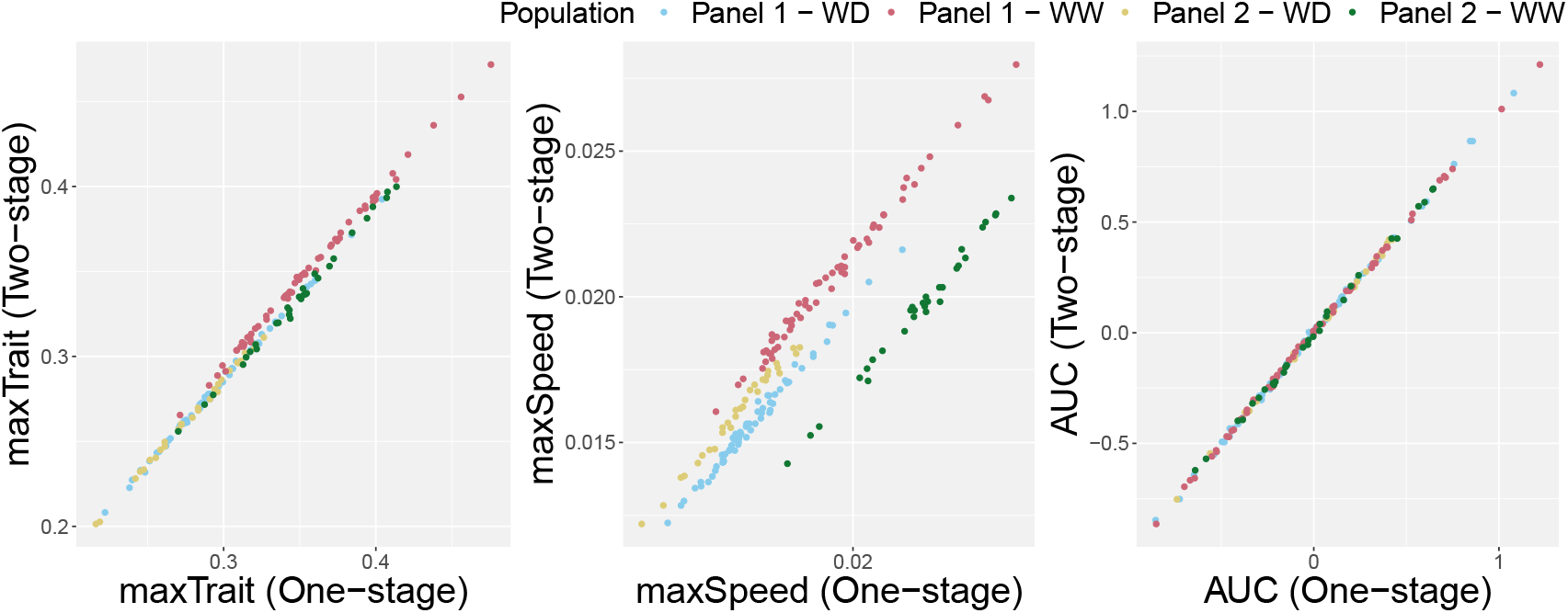
Results for the PhenoArch platform: Bivariate scatterplots with the extracted attributes at the genotype level. Each scatterplot depicts the comparison between the one- and the two-stage approaches for one feature. Colours represent the populations (panel-by-water regime combination).

## 3 Application

### 3.1 Data description

We illustrate the performance of our proposal with data from the PhenoArch platform (INRAE, Montpellier; Cabrera-Bosquet et al., 2016). This dataset was also used to illustrate the results of the two-stage approach proposed by Pérez-Valencia et al. (2022). We briefly describe the details needed to understand the application here. The platform comprises 1680 pots (with single plants in pots) on a rectangular grid of *R* = 60 rows and *C* = 28 columns. The experiment included two genotype panels (Panel 1 and Panel 2) of commercial maize hybrids representative of breeding history in Europe and US during the last 60 years. A total of 90 genotypes were tested: 60 in Panel 1 and 30 in Panel 2. Each genotype was replicated between 4 (Panel 2) and 14 (Panel 1) times. All genotypes were tested under two levels of soil water content: WD - mild water deficit (soil water potential of —0.5 MPa) and WW - retention capacity (soil water potential of —0.05 MPa). The experiment was conducted in mid-spring 2017 between April 13th and May 13th, i.e., between 103 and 133 days since January 1st, (hereafter DOY, Day of the Year). The trait of interest is the leaf area (*m*^2^ plant^−1^), measured on *M* = 1648 plants at *n* = 31 timepoints. As in Pírez-Valencia et al. (2022), we do not consider the panel by water regime crossed-effects structure, instead, the two panels and the two water regimes are combined, such that we have *K* = 4 “populations” (Panel-by-water regime combination: Panel 1 - WD with 589 plants, Panel 1 - WW with 823 plants, Panel 2 - WD with 118 plants and Panel 2 - WW with 118 plants), and a total of *L* = 180 “genotypes” (60 genotypes in Panel 1 - WD, 60 genotypes in Panel 1 - WW, 30 genotypes in Panel 2 - WD and 30 genotypes in Panel 2 - WW). Figure 3 depicts the evolution over time of the raw leaf area for plants in each panel-by-water regime combination. A large amount of missing values characterises this dataset. A total of 37970 observations are available (out of 51088 = 1648 plants × 31 times).

### Results: One vs two-stage approach

The aim of this section is twofold. We present the results by modelling the PhenoArch dataset using the one-stage approach, and at the same time, we compare these results with the ones obtained with the two-stage approach. For the one-stage approach, we fit model (3) with *b*_row_ = *b*_col_ = 11 B-splines for the random row, *f_r_*, and column, *f_c_*, effects, *b_r_* = *b_c_* = *b_t_* = 8 for the spatio-temporal smooth function, *f*_ST_, and *b*_pop_ = *b*_geno_ = *b*_plant_ = 11 for the hierarchical components, *f_p_*, *f_g_* and *f_i_*. Under this configuration, the mixed model formulation of the one-stage approach has a total of 21624 regression coefficients (both fixed and random) and 20 variance parameters. As for the simulation experiment, to make the one- and two-stage approaches comparable, for the two-stage approach we set the dimensions of the B-spline basis that are common to both approaches to the same values. Therefore, the mixed model formulation in the second-stage of the two-stage approach has a total of 20152 regression coefficients (both fixed and random) and 11 variance parameters. Computations were performed in a (64-bit) R 4.2.1 and a 1.60GHz Dual-Core^TM^ i5 processor computer with 16GB of RAM and macOS Monterrey Version 12.5. Estimation took approximately one and a half minutes for the two-stage approach (20 seconds the first stage and 60 seconds the second stage) and 25 minutes for the one-stage approach. We highlight here that the computational time of the two-stage approach reported in Pérez-Valencia et al. (2022) has been improved by building the mixed model equations in a more efficient way and integrating the methods from the `R`-package `LMMsolver` in the codes to estimate the variance parameters.

In Figure 4, we first illustrate the spatial trend obtained with both approaches at four different time points. As expected, the spatial trends estimated by the one-stage approach (Figure 4(a)), show that they vary smoothly over time. In contrast, the spatial trends obtained with the two-stage approach (Figure 4(b)), exhibit more marked differences among time measurements because analyses in the first-stage are performed separately per time point. Consequently, information on spatial heterogeneity is not shared across different measurement times. Finally, a detailed look at the scale of these two plots reveals a small spatial effect for this particular dataset.

For the two approaches, we also obtained (as for the simulation study) estimated curves at the three-level hierarchy. We focus here on the results at the genotype level, which is the decision-making level for plant breeders and the level at which we can study genotype-by-environment interactions. In Figure 5, we use four genotypes per panel to illustrate our results for the one-stage (continuous lines) and two-stage (dotted lines) approaches. Genotypes were chosen such that two of them have the best and worst performance and the other two have an intermediate performance (results for all genotypes can be found in Web Figure 7). Figure 5(a) shows different growth patterns under both water regimes, e.g., well watered (WW) plants grow faster than plants with water deficit (WD) for both panels. Small differences are observed between the one and two-stage approaches, being “Panel 1 - WW” the population with the most considerable difference. Figure 5(b) depicts the first-order derivative for the genotype-specific growth curves. These curves show differences in the speed of leaf area growth (or leaf area growth rate) for the selected genotypes. From these curves, information about the maximum slope and the time at which this maximum is reached are important for understanding plant development. For instance, if the first maximum point is analysed in both panels, growth rates for WW plants are higher than those for WD, but WD plants reach the maximum point earlier than WW. We have found that the major discrepancy between the two approaches correspond to the estimated first-order derivative curves. This is in concordance with the results of the simulation study and the analysis of other HTP datasets. Finally, genotype-specific deviations in Figure 5(c) allow evaluating differences in genotypic performance. That is, positive and negative deviations refer to better and worse genotypic performance compared to the mean population trajectory.

For instance, in Panel 1 and under both water regimes, genotype 48 performs better than genotype 29. This figure also allows us to analyse the genotype-by-water regime interaction. Note that genotypes 43 (in blue) and 44 (in purple) in Panel 1 have similar performance under WD, but their performances differ under WW. For Panel 2, we see that the curves for genotypes 03 (in red) and 15 (in brown) differ under WD, but become similar under WW. Differences between both approaches for these curves are minimal.

One of the most important aspects in these analyses is whether decision-making changes with the approach used. To address this question, we extracted some time-independent features from the estimated curves to characterise genotypes. We are aware that this data does not have information for the stationary phase common in the classic growth curve analysis. Consequently, the feature extraction is limited by the time window at which plants were measured. However, we calculate three features for all genotypes: (1) the maximum corrected leaf area (maxTrait) from the estimated genotype-specific growth curves (Figure 5(a)), (2) the maximum speed rate (maxSpeed), before *t* = 130, from the estimated first-order derivatives for the genotype-specific growth curves (Figure 5(b)), and (3) the area under the estimated genotype-specific deviations (AUC; Figure 5(c)). The two approaches are compared by bivariate scatter plots for each feature, as shown in Figure 6. The maxTrait and the AUC are the strongest correlated features, indicating small differences between both approaches for the decision-making process. As expected and in concordance with the simulation study, maxSpeed is the most sensitive feature to differences between both approaches (the results under the WW treatment show, for both panels, the highest difference). However, the values of maxSpeed obtained using both approaches also show a high correlation within a panel-by-water regime combination. In brief, results of both the simulation study and the application are consistent. That is, slight differences are detected when using the one- and two-stage approaches for most situations, with the estimation of the first-order derivative being the result with the most noticeable differences between the approaches used.

## 4 Discussion

Modelling large spatio-temporal and hierarchical data, as obtained from HTP platforms, has become a challenge for statisticians and plant breeding practitioners. One-stage approaches represent an elegant solution that overcomes the loss of information given by stage-wise approaches, but the computational complexity still persists (Verbyla et al., 2021). To tackle this problem, in this paper, we have proposed a one-stage spatio-temporal P-spline-based hierarchical approach for the modelling of the genetic and non-genetic variation of HTP data. To make our proposal computationally affordable, we combine different specialised methods that take advantage of the sparse model matrices structure (Boer & van Rossum, 2022), the array data structure (Currie et al., 2006), and the non-standard form of the variance-covariance matrix (Rodríguez-Álvarez et al., 2019; Rodríguez-Álvarez et al., 2015).

To assess the performance of our method and compare its results with the two-stage approach proposed by Pérez-Valencia et al. (2022), we designed a simulation study. The simulated spatio-temporal and hierarchical data structure that we proposed allowed us to reproduce data common to HTP experiments while evaluating results at higher levels of the hierarchy (e.g. population and genotype levels). We found that for most simulated situations, there was no clear difference between the two approaches, except when non-nested B-spline bases were used. The more considerable differences were detected for the first-order derivative curves, where the one-stage approach outperforms the two-stage approach, and the smaller the bases, the better the performance for both approaches. We also illustrated our approach with data from the PhenoArch platform. The results at the genotype level follow the simulation results, i.e., minor differences between the two approaches were detected, except for the first-order derivatives for the genotype-specific growth curves. Nevertheless, highly correlated results were obtained between the two approaches when we extracted important time-independent features from these curves for, e.g., genotype selection.

We offer a good starting point (“proof-of-concept”) to analyse spatio-temporal and hierarchical HTP data. We are aware that some limitations have to be addressed. We have proposed a P-spline-based approach, which means that as the number of plants and basis dimensions increase, so does the computation time (scalability problem). Nevertheless, for the PhenoArch example, it only took 25 minutes to obtain the results without any convergence issues. We recommend using the same number of B-spline basis functions for the three levels of the hierarchy, as well as for the row and column random effects. Regarding the number of B-spline basis functions used for the three-dimensional surface (in the row, column and time directions), we suggest keeping them relatively small to enable the solution to run on standard computers. Although we present a general approach, our implementation (code) is very specific. For instance, we only consider the spatio-temporal smooth function and random effects for rows and columns as non-genetic effects, but no other experimental factors were taken into account. Furthermore, we only consider a nested structure in the data, but, e.g., the PhenoArch example has a crossed-effect structure, where explicitly modelling the genotype-by-treatment interaction would be more appropriate.

In our experience, it is computationally simpler to obtain results from stage-wise approaches. However, one-stage approaches will always represent statistical adequacy. For instance, in our work with the analysis of different HTP data, we have observed that the one-stage approach performs better in the presence of missing data (since it borrows strength across plant curves; more exploration in this direction is required using simulated data). We believe a good practice would be to use a two-stage approach as a starting point to establish the basis for a model in one stage. For this purpose, we make publicly available the `R`-functions with the one- and two-stage approaches implementation at https://gitlab.bcamath.org/dperez/htp_one_stage_approach, where we also provide the code and data to reproduce the analyses and results of the application shown in this paper. `R`-functions to analyse data with the two-stage approach are already freely available in the statgenHTP `R`-package (https://CRAN.R-project.org/package=statgenHTP, Millet et al., 2022). We expect soon to add the one-stage approach functions to the aforementioned package to simplify the access.

## Acknowledgements

This research is supported by the Basque Government through the BERC 2022-2025 program and by the Ministry of Science and Innovation: BCAM Severo Ochoa accreditation CEX2021-001142-S/MICIN/AEI/10.13039/501100011033, by the Ramon y Cajal Grant RYC2019-027534-I, and by EU H2020 grant agreement ID 731013 (EPPN^2020^). We thank Llorenç Cabrera-Bosquet and François Tardieu for sharing with us the PhenoArch data, and Bart-Jan van Rossum for helping us to integrate and improve the `R`-functions to fit our proposal.

## Data availability

The PhenoArch dataset, analysed during the current study, is available with the paper and its “Supporting information”.

## Supporting information

Supporting information (code, data and additional figures) is available for this paper in the Basque Center for Applied Mathematics repository, https://gitlab.bcamath.org/dperez/htp_one_stage_approach.

