## Supporting information for "A one-stage approach for the spatio-temporal analysis of high-throughput phenotyping data"

This document contains additional information to the paper "A one-stage approach for the spatio-temporal analysis of high-throughput phenotyping data". Web Appendix A briefly describes the P-splines formulation in one, two and three dimensions, and their parameterisation in the linear mixed model form. Additional figures for the simulation study of Section 2.3 and for the application of Section 3 in the main text are shown in Web Appendix B and Web Appendix C, respectively.

#### Web Appendix A Some notes on P-splines and mixed models

In this section we provide some background on P-splines as well as their linear mixed model form. We start with the univariate case and then move to the three-dimensional case.

Let $\boldsymbol{y} = (y_1, \ldots, y_n)^T$ be a vector of $n$ observations, and consider the model

$$y_i = f(t_i) + \varepsilon_i,\ \varepsilon_i \sim N(0, \sigma^2),\ i = 1, \ldots, n,$$

where $f$ is a smooth and unknown function to be estimated from the data. In P-splines, the function $f$ is approximated by a linear combination of $b_f$ B-spline basis functions, i.e., $f(t) = \sum_{j=1}^{b_f} \theta_j B_j(t)$. In matrix form, we write

$$\boldsymbol{f} = \boldsymbol{B}\boldsymbol{\theta}, \tag{A1}$$

where $\boldsymbol{f} = (f(t_1), \ldots, f(t_n))^T$, $(\boldsymbol{B})_{lj} = B_j(t_l)$, and $\boldsymbol{\theta} = \left(\theta_1, \ldots, \theta_{b_f}\right)^T$. Smoothness is ensured by penalising the differences of order $q$ of coefficients associated with adjacent B-spline basis functions (in this work we use second-order difference penalties, i.e, $q = 2$). In particular, the penalty takes the following form

$$\lambda \boldsymbol{\theta}^T \boldsymbol{D}^T \boldsymbol{D} \boldsymbol{\theta}, \tag{A2}$$

where $\boldsymbol{D}$ is a matrix that forms differences of order $q$, and $\lambda$ is the smoothing parameter (it sets the weight of the penalty: the larger the value of $\lambda$, the smoother the result will be).

We now provide the linear mixed model form of (A1). Here, the design matrix $\boldsymbol{B}$ and the vector of regression coefficients $\boldsymbol{\theta}$ are reformulated in such a way that

$$\boldsymbol{f} = \boldsymbol{B}\boldsymbol{\theta} = \boldsymbol{X}\boldsymbol{\beta} + \boldsymbol{Z}\boldsymbol{u}, \text{ with } \boldsymbol{u} \sim N(\boldsymbol{0}, \boldsymbol{G}).$$

where $\boldsymbol{X}$ and $\boldsymbol{Z}$ are the mixed model design matrices, and $\boldsymbol{\beta}$ and $\boldsymbol{u}$ are the vector of fixed and random effect coefficients, respectively. There are different ways in which $\boldsymbol{X}$ and $\boldsymbol{Z}$ can be obtained (Currie & Durban, 2002; Currie et al., 2006; Lee & Durban, 2011; Wand, 2003). We follow Currie et al. (2006) and Lee and Durban (2011), and use the singular value decomposition of the penalty matrix $\boldsymbol{P} = \boldsymbol{D}^T\boldsymbol{D}$. In particular, let $\boldsymbol{P} = \boldsymbol{U}\boldsymbol{\Lambda}\boldsymbol{U}^T$, be the eigenvalue decomposition of $\boldsymbol{P}$, where $\boldsymbol{U}$ is the matrix of eigenvectors and $\boldsymbol{\Lambda}$ is the diagonal matrix eigenvalues. Let us also denote $\boldsymbol{U}_+$ ($\boldsymbol{\Lambda}_+$) and $\boldsymbol{U}_0$ ($\boldsymbol{\Lambda}_0$) the sub-matrices corresponding to the non-zero and zero eigenvalues, respectively. We note that for second-order difference penalties there are two zero eigenvalues, and thus $\boldsymbol{\Lambda}_0$ is a 2-by-2 matrix of zeroes. It is easy to show that (A1) can then be reparameterised as

$$\boldsymbol{f} = \boldsymbol{B}\boldsymbol{\theta} = \boldsymbol{B}\boldsymbol{U}\boldsymbol{U}^T\boldsymbol{\theta} = \boldsymbol{X}\boldsymbol{\beta} + \boldsymbol{Z}\boldsymbol{u},$$

where $\boldsymbol{X} = \boldsymbol{B}\boldsymbol{U}_0$, $\boldsymbol{Z} = \boldsymbol{B}\boldsymbol{U}_+$, $\boldsymbol{\beta} = \boldsymbol{U}_0^T\boldsymbol{\theta}$, and $\boldsymbol{u} = \boldsymbol{U}_+^T\boldsymbol{\theta}$. Furthermore, $\boldsymbol{\theta} = \boldsymbol{U}_0\boldsymbol{\beta} + \boldsymbol{U}_+\boldsymbol{u}$, and thus the penalty (A2) can be rewritten as

$$\lambda \boldsymbol{\theta}^T \boldsymbol{P} \boldsymbol{\theta} = \lambda \boldsymbol{u}^T \boldsymbol{\Lambda}_+ \boldsymbol{u}.$$

That is to say, only the vector $\boldsymbol{u}$ is penalised, while $\boldsymbol{\beta}$ it is not. In the equivalent mixed model, it implies that $\boldsymbol{\beta}$ is a vector of fixed effect coefficients, and that $\boldsymbol{u}$ is a vector of random effect coefficients, assumed to be distributed according to a multivariate Gaussian with zero mean and precision matrix (the inverse of the variance-covariance matrix) given by $\boldsymbol{G}^{-1} = \sigma_f^{-2}\boldsymbol{\Lambda}_+$, where $\sigma_f^2 = \sigma^2/\lambda$ (i.e., $\boldsymbol{G} = \sigma_f^2\boldsymbol{\Sigma}$, with $\boldsymbol{\Sigma} = \boldsymbol{\Lambda}_+^{-1}$, is the variance-covariance matrix). To finish this part, we note that based on the eigenvalue decomposition we have that $\boldsymbol{X} = \boldsymbol{B}\boldsymbol{U}_0$. However, it is sometimes more convenient (or, at least, it helps understanding the unpenalised/fixed part of a P-spline) to take $\boldsymbol{X} = [\boldsymbol{1}_n \mid \boldsymbol{t}]$. In other words, when using P-splines in combination with a second-order penalty, the space of functions that are not penalised corresponds to the polynomials of degree 1. Another possible specification for $\boldsymbol{X}$ is the one proposed by Wood et al. (2013), in which we also obtain a design matrix with a constant column (not necessarily of ones). For this purpose, we first obtain $\tilde{\boldsymbol{X}} = \boldsymbol{B}\boldsymbol{U}_0$ as before, and then compute the singular value decomposition of $\boldsymbol{F}^T\boldsymbol{F} = \boldsymbol{V}\boldsymbol{\Omega}\boldsymbol{V}^T$, with $\boldsymbol{F} = \tilde{\boldsymbol{X}} - \boldsymbol{1}\boldsymbol{1}^T\tilde{\boldsymbol{X}}/n$ (i.e., based on centered values for $\tilde{\boldsymbol{X}}$), and define $\boldsymbol{X} = \boldsymbol{B}\boldsymbol{U}_0\boldsymbol{V}$. In our experience, we have found this approach to be numerically more stable and it is the one we implement in our code.

When it comes to extend the P-spline principles to the three-dimensional case, we first approximate the three-dimensional smooth function we are interested in estimating by using the tensor-product of three marginal B-splines basis, i.e.,

$$f_{\mathrm{ST}}(r, c, t) = \sum_{j=1}^{b_r}\sum_{q=1}^{b_c}\sum_{v=1}^{b_t} \theta_{jqv} B_{1j}(r) B_{2q}(c) B_{3v}(t),$$

and smoothness is achieved by penalising coefficient differences along each covariate (in our case, row number, column number and time), i.e., the (anisotropic) penalty in three dimensions is

$$\boldsymbol{\theta}_{\mathrm{ST}}^{\top}\left(\lambda_1\left(\boldsymbol{D}_1^{\top}\boldsymbol{D}_1 \otimes \boldsymbol{I}_{b_c} \otimes \boldsymbol{I}_{b_t}\right) + \lambda_2\left(\boldsymbol{I}_{b_r} \otimes \boldsymbol{D}_2^{\top}\boldsymbol{D}_2 \otimes \boldsymbol{I}_{b_t}\right) + \lambda_3\left(\boldsymbol{I}_{b_r} \otimes \boldsymbol{I}_{b_c} \otimes \boldsymbol{D}_3^{\top}\boldsymbol{D}_3\right)\right)\boldsymbol{\theta}_{\mathrm{ST}}, \tag{A3}$$

where $\boldsymbol{\theta}_{\mathrm{ST}} = (\theta_{111}, \ldots, \theta_{11b_t}, \theta_{121}, \ldots, \theta_{12b_t}, \ldots, \theta_{1b_c1}, \ldots, \theta_{1b_cb_t}, \theta_{211}, \ldots, \theta_{21b_t}, \ldots, \theta_{b_rb_c1}, \ldots, \theta_{b_rb_cb_t})^T$, $\boldsymbol{D}_m$ $(m = 1, 2, 3)$ are difference matrices of order 2, and $\lambda_1$, $\lambda_2$ and $\lambda_3$ are the smoothing parameters (Eilers & Marx, 2003). Using the notation in Section 2 of the main text, in matrix form we write

$$f_{\mathrm{ST}}(\boldsymbol{r}, \boldsymbol{c}, \boldsymbol{t}) = \left(\left(\boldsymbol{B}_1 \square \boldsymbol{B}_2\right) \otimes \boldsymbol{B}_3\right)\boldsymbol{\theta}_{\mathrm{ST}}, \tag{A4}$$

where $(\boldsymbol{B}_1)_{ij} = B_{1j}(r(i))$, $(\boldsymbol{B}_2)_{iq} = B_{2q}(c(i))$ and $(\boldsymbol{B}_3)_{lv} = B_{3v}(t_l)$. To obtain the linear mixed model form of (A4), we follow Lee and Durban (2011). As shown in that paper, in this case the mixed model design matrices are

$$\begin{aligned}
\boldsymbol{X}_{\mathrm{ST}} &= [(\boldsymbol{X}_r \square \boldsymbol{X}_c) \otimes \boldsymbol{X}_t],\\
\boldsymbol{Z}_{\mathrm{ST}} &= [(\boldsymbol{Z}_r \square \boldsymbol{X}_c) \otimes \boldsymbol{X}_t \mid (\boldsymbol{X}_r \square \boldsymbol{Z}_c) \otimes \boldsymbol{X}_t \mid (\boldsymbol{X}_r \square \boldsymbol{X}_c) \otimes \boldsymbol{Z}_t \mid \\
&\qquad (\boldsymbol{Z}_r \square \boldsymbol{Z}_c) \otimes \boldsymbol{X}_t \mid (\boldsymbol{Z}_r \square \boldsymbol{X}_c) \otimes \boldsymbol{Z}_t \mid (\boldsymbol{X}_r \square \boldsymbol{Z}_c) \otimes \boldsymbol{Z}_t \mid (\boldsymbol{Z}_r \square \boldsymbol{Z}_c) \otimes \boldsymbol{Z}_t],
\end{aligned} \tag{A5}$$

where $\boldsymbol{X}_r = \boldsymbol{B}_1\boldsymbol{U}_{10}$, $\boldsymbol{Z}_r = \boldsymbol{B}_1\boldsymbol{U}_{1+}$, $\boldsymbol{X}_c = \boldsymbol{B}_2\boldsymbol{U}_{20}$, $\boldsymbol{Z}_c = \boldsymbol{B}_2\boldsymbol{U}_{2+}$, $\boldsymbol{X}_t = \boldsymbol{B}_3\boldsymbol{U}_{30}$, $\boldsymbol{Z}_t = \boldsymbol{B}_3\boldsymbol{U}_{3+}$, and the precision matrix for the vector of random effects is

$$\begin{aligned}
\boldsymbol{G}_{\mathrm{ST}}^{-1} = \mathrm{blockdiag}\Bigg(&\frac{1}{\sigma_1^2}\boldsymbol{\Lambda}_{1+} \otimes \boldsymbol{I}_2 \otimes \boldsymbol{I}_2, \frac{1}{\sigma_2^2}\boldsymbol{I}_2 \otimes \boldsymbol{\Lambda}_{2+} \otimes \boldsymbol{I}_2, \frac{1}{\sigma_3^2}\boldsymbol{I}_2 \otimes \boldsymbol{I}_2 \otimes \boldsymbol{\Lambda}_{3+},\\
&\frac{1}{\sigma_1^2}\boldsymbol{\Lambda}_{1+} \otimes \boldsymbol{I}_{b_c-2} \otimes \boldsymbol{I}_2 + \frac{1}{\sigma_2^2}\boldsymbol{I}_{b_r-2} \otimes \boldsymbol{\Lambda}_{2+} \otimes \boldsymbol{I}_2,\\
&\frac{1}{\sigma_1^2}\boldsymbol{\Lambda}_{1+} \otimes \boldsymbol{I}_2 \otimes \boldsymbol{I}_{b_t-2} + \frac{1}{\sigma_3^2}\boldsymbol{I}_{b_r-2} \otimes \boldsymbol{I}_2 \otimes \boldsymbol{\Lambda}_{3+},\\
&\frac{1}{\sigma_2^2}\boldsymbol{I}_2 \otimes \boldsymbol{\Lambda}_{2+} \otimes \boldsymbol{I}_{b_t-2} + \frac{1}{\sigma_3^2}\boldsymbol{I}_2 \otimes \boldsymbol{I}_{b_c-2} \otimes \boldsymbol{\Lambda}_{3+},\\
&\frac{1}{\sigma_1^2}\boldsymbol{\Lambda}_{1+} \otimes \boldsymbol{I}_{b_c-2} \otimes \boldsymbol{I}_{b_t-2} + \frac{1}{\sigma_2^2}\boldsymbol{I}_{b_r-2} \otimes \boldsymbol{\Lambda}_{2+} \otimes \boldsymbol{I}_{b_t-2} + \frac{1}{\sigma_3^2}\boldsymbol{I}_{b_r-2} \otimes \boldsymbol{I}_{b_c-2} \otimes \boldsymbol{\Lambda}_{3+}\Bigg),
\end{aligned}$$

where $\sigma_1^2 = \sigma^2/\lambda_1$, $\sigma_2^2 = \sigma^2/\lambda_2$ and $\sigma_3^2 = \sigma^2/\lambda_3$. In the above expressions, $\boldsymbol{U}_{m+}$ ($\boldsymbol{\Lambda}_{m+}$) and $\boldsymbol{U}_{m0}$ ($\boldsymbol{\Lambda}_{m0}$) are the sub-matrices corresponding, respectively, to the non-zero and zero eigenvalues of $\boldsymbol{D}_m^\top \boldsymbol{D}_m$ ($m = 1, 2, 3$; see (A3)). By replacing in (A5) $\boldsymbol{X}_r = \boldsymbol{B}_1\boldsymbol{U}_{10}$, $\boldsymbol{X}_c = \boldsymbol{B}_2\boldsymbol{U}_{20}$ and $\boldsymbol{X}_t = \boldsymbol{B}_3\boldsymbol{U}_{30}$ by, respectively, $\boldsymbol{X}_r = [\boldsymbol{1}_M \mid \boldsymbol{r}]$, $\boldsymbol{X}_c = [\boldsymbol{1}_M \mid \boldsymbol{c}]$ and $\boldsymbol{X}_t = [\boldsymbol{1}_n \mid \boldsymbol{t}]$, (or more precisely, by the matrices obtained using Wood et al. (2013) approach, see above) and after some reorganisation of the matrices as well as the removal of the redundant components, we arrive to the expression given in equations (6) and (7) of the main text (and the ANOVA-type decomposition presented in (5))

### Web Appendix B Additional results for the simulation study

In this section we present additional information for the simulation study presented in Section 2.3 of the main text. Web Appendix B.1 describes the steps followed for the generation of the simulated data. Web Appendix B.2 shows all the additional figures and tables.

#### Web Appendix B.1 Steps for the generation of the simulated data

Using the notation in Section 2.3.1 of the main text, the spatio-temporal data are generated from the population to the plant level in the following five steps

**Step 1.** Generate one population trajectory, $\boldsymbol{f}_p = (f_p(t_1), \ldots, f_p(t_n))^T$, from the growth logistic curve model. Following the notation in Li and Sillanpää (2015), we consider $\boldsymbol{f}_p = a/(1 + e^{c(b-\boldsymbol{t})})$, where $a$ is the asymptote, $b$ is the inflection point, and $c$ is the growth rate. Additionally, it can be shown that the first-order derivative of this function with respect to time, $t$, is $\boldsymbol{f}'_p = ace^{c(b-\boldsymbol{t})}/(1 + e^{c(b-\boldsymbol{t})})^2$.

**Step 2.** Generate $L$ genotype-specific deviations, $\boldsymbol{f}_g = (f_g(t_1), \ldots, f_g(t_n))^T \overset{\text{iid}}{\sim} N(\boldsymbol{0}, \boldsymbol{\Sigma}_{\text{gen}})$ ($g = 1, \ldots, L$). For the $n \times n$ variance-covariance matrix $\boldsymbol{\Sigma}_{\text{gen}}$, we consider a first-order autoregressive structure with heterogeneous variance (ARH(1), with a slight modification to the structure presented by Wolfinger, 1996, to account for variance increasing with time structure), that is $(\boldsymbol{\Sigma}_{\text{gen}})_{jk} = (1/(1-\rho^2))s_{jk}^{\text{gen}}\rho^{d_{jk}}$, where $\rho$ is the autocorrelation parameter, $d_{jk}$ is the euclidean distance between time points $t_j$ and $t_k$ (i.e., $d_{jk} = |t_k - t_j|$)), and $s_{jk}^{\text{gen}} = \sigma_{\text{gen}}^4 h(t_j)h(t_k)$ are the elements of the heterogeneous variance-covariance matrix. Here, $\sigma_{\text{gen}}^2$ is the between genotype (deviation) variability, and $h(\cdot)$ is a function that is quadratic in $t$. The genotype trajectories can be obtained as the sum of the population trajectory and the genotype deviations, i.e., $\boldsymbol{f}_p + \boldsymbol{f}_{g(p)}$, as depicted in Figure 1 of the main text.

**Step 3.** Generate $M$ $(= R \times C)$ plant-specific deviations, $\boldsymbol{f}_i = (f_i(t_1), \ldots, f_i(t_n))^T \overset{\text{iid}}{\sim} N(\boldsymbol{0}, \boldsymbol{\Sigma}_{\text{plant}})$ ($i = 1, \ldots, M$). We follow the same ideas used in **Step 2** to obtain the $n \times n$ variance-covariance matrix $\boldsymbol{\Sigma}_{\text{plant}}$. We use $\sigma_{\text{plant}}^2$ for the between plants (deviation) variability, and $s_{jk}^{\text{plant}} = \sigma_{\text{plant}}^4 h(t_j)h(t_k)$ for the elements of the heterogeneous variance-covariance matrix.

**Step 4.** Generate $M = R \times C$ spatio-temporal correlated noise curves, $\boldsymbol{\varepsilon}(\boldsymbol{r}, \boldsymbol{c}, \boldsymbol{t}) = (\varepsilon(r(1), c(1), \boldsymbol{t}), \varepsilon(r(2), c(2), \boldsymbol{t}), \ldots, \varepsilon(r(M), c(M), \boldsymbol{t}))^T \sim N(\mathbf{0}, \boldsymbol{\Sigma}_\varepsilon)$, where $\varepsilon(r(i), c(i), \boldsymbol{t}) = (\varepsilon(r(i), c(i), t_1), \ldots, \varepsilon(r(i), c(i), t_n))$ is the noise curve for the $i$th plant. For the $(RCn) \times (RCn)$ variance-covariance matrix $\boldsymbol{\Sigma}_\varepsilon$, we use a space-time separable covariance model $\boldsymbol{\Sigma}_\varepsilon = \boldsymbol{\Sigma}_{\mathrm{S}} \otimes \boldsymbol{\Sigma}_{\mathrm{T}}$, where $\boldsymbol{\Sigma}_{\mathrm{S}}$ is a $(RC) \times (RC)$ isotropic and homogeneous spatial variance-covariance matrix (Matérn, Guttorp & Gneiting, 2006), and $\boldsymbol{\Sigma}_{\mathrm{T}}$ is a $n \times n$ temporal ARH(1) variance-covariance matrix. In particular,

$$(\boldsymbol{\Sigma}_{\mathrm{S}})_{jk} = \frac{1}{2^{\omega-1}\Gamma(\omega)} \left( \frac{s^{\mathrm{S}}_{jk}}{\nu} \right)^{\omega} \kappa_\omega \left( \frac{s^{\mathrm{S}}_{jk}}{\nu} \right), \quad (\boldsymbol{\Sigma}_{\mathrm{T}})_{jk} = \frac{1}{1 - \rho_\varepsilon^2} s^{\varepsilon}_{jk} \rho_\varepsilon^{d_{jk}},$$

where $\kappa_\omega(\cdot)$ denotes the modified Bessel function of the third kind and order $\omega$, with $\omega > 0$ being a smoothness parameter, $\Gamma(\cdot)$ is the Gamma function, $\nu > 0$ is the scale parameter of the correlation function, and $s^{\mathrm{S}}_{jk} = \sqrt{(c(k) - c(j))^2 + (r(k) - r(j))^2}$ is the euclidean distance between two plants locations, $(r(k), c(k))$ and $(r(j), c(j))$. As for **Steps 2** and **3**, $s^{\varepsilon}_{jk} = \sigma_\varepsilon^4 h_\varepsilon(t_j) h_\varepsilon(t_k)$, where $\sigma_\varepsilon^2$ is the residual variance, $h_\varepsilon(\cdot)$ is a function of $t$ to the power of 0.8 and $\rho_\epsilon$ is the autocorrelation parameter.

In this step, genotypes are assigned to spatial positions following a randomised complete block design, such that each replicate (plant) of a genotype is present in just one block. Depending on the number of replicates ($m_g$), blocks are accommodated in the row or column direction such that the size of the blocks is the ratio between the number of rows (or columns) and the number of replicates (see Web Figure 1 for an illustrative example of the randomisation).

**Step 5.** Calculate $M$ plant trajectories as the sum of the population trajectories, the genotype and plant deviations and the spatio-temporal correlated noise, i.e., $\boldsymbol{y}_i = \boldsymbol{f}_p + \boldsymbol{f}_{g(i)} + \boldsymbol{f}_i + \boldsymbol{\varepsilon}(r(i), c(i), \boldsymbol{t})$.

#### Web Appendix B.2 Additional figures and tables for the simulation study

Web Figure 2 depicts the heritability over time under the eight simulation scenarios. Web Table 1 presents the simulation settings used to generate the spatio-temporal data. To assess the performance of the one-stage approach, we propose to compare five different configurations for the dimensions of the B-spline bases for the hierarchical components, $f_p$, $f_g$ and $f_i$. One of the configurations refers to a non-nested bases; Web Figure 3 depicts an illustrative visualisation for this concept. Finally, Web Figures 4, 5 and 6 aim to support the simulation results (for six of the eight simulation scenarios reported).

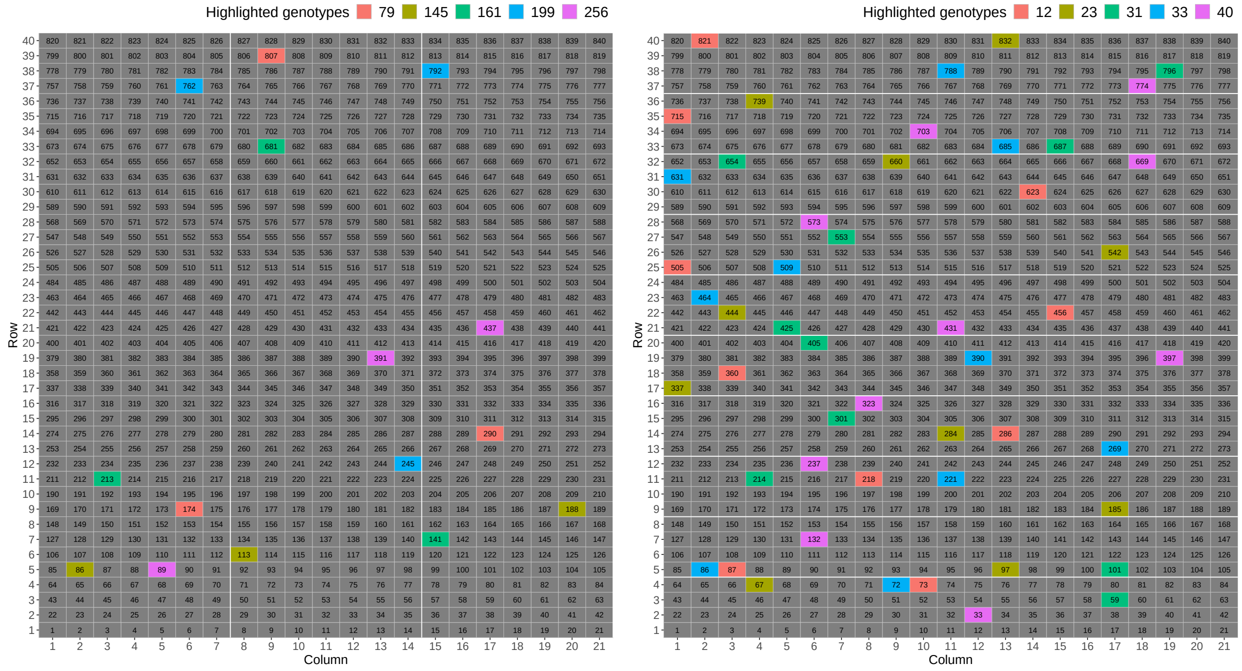

(a) $m_g = 3$ (b) $m_g = 10$

Web Figure 1: Illustrative visualisation of the grid with the randomisation used for two simulated datasets (as illustration) with (a) $m_g = 3$, and (b) $m_g = 10$ replicates per genotype. The size of the grid is $R \times C = 40 \times 21$, for a total of $M = 840$ plants. Each cell represents a plant ($i = 1, \ldots, M$), which is identified by its row and column position, i.e., the $i$th plant has coordinates $(r(i), c(i))$ (e.g., plant $i = 457$ has position $(r(457), c(457)) = (22, 16)$). Colours depict replicates in five selected genotypes (as illustration).

| Level | Description | Parameter | Value |
|---|---|---|---|
| Experimental design | Number of populations | $p$ | 1 |
| | Number of replicates per genotype | $m_g$ | 3 and 10 |
| | Number of timepoints | $n$ | 25 |
| | Number of rows | $R$ | 40 |
| | Number of columns | $C$ | 21 |
| | Number of genotypes | $L$ | $RC/m_g$ |
| | Number of plants | $M$ | $Lm_g$ |
| Population level | Asymptote | $a$ | 0.6 |
| | Inflection point | $b$ | 12.5 |
| | Growth rate | $c$ | 0.4 |
| Genotype level | Between genotypes (deviations) variability | $\sigma^2_{\text{gen}}$ | $\sigma^2_l = 8 \times 10^{-7}$ and $\sigma^2_h = 1.2 \times 10^{-6}$ |
| | Time function | $h(t)$ | $8 \times 10^{-1} t^2$ |
| | Autocorrelation | $\rho$ | 0.9999 |
| Plant level | Between plants (deviations) variability | $\sigma^2_{\text{plant}}$ | $\sigma^2_l = 8 \times 10^{-7}$ and $\sigma^2_h = 1.2 \times 10^{-6}$ |
| | Time function | $h(t)$ | $8 \times 10^{-1} t^2$ |
| | Autocorrelation | $\rho$ | 0.9999 |
| Spatio-temporal correlated noise | Residual variance | $\sigma^2_\varepsilon$ | $1 \times 10^{-8}$ |
| | Time function | $h_\varepsilon(t)$ | $1000 \times t^{0.8}$ |
| | Autocorrelation | $\rho_\varepsilon$ | 0.5 |
| | Smoothness parameter | $\omega$ | 10000 |
| | Scale parameter | $\nu$ | 5 |

Web Table 1: Simulation settings. Reference values are based on the leaf area ($m^2$ plant$^{-1}$) data from the PhenoArch platform. For this data, the maximum leaf area is approximately 0.5 (we fixed the asymptote for the population trajectory at $a = 0.6$), the average growth rate is 0.01 (we increased this value to $c = 0.4$ to obtain S-shape curves), the between plants variability at the beginning of the experiment is $4.5 \times 10^{-7}$ (we set $\sigma^2_l = 8 \times 10^{-7}$ and $\sigma^2_h = 1.2 \times 10^{-6}$), and the maximum first-order autocorrelation for observations of plant trajectories is 0.9986 (we used $\rho = 0.9999$ to obtain smoother curves).

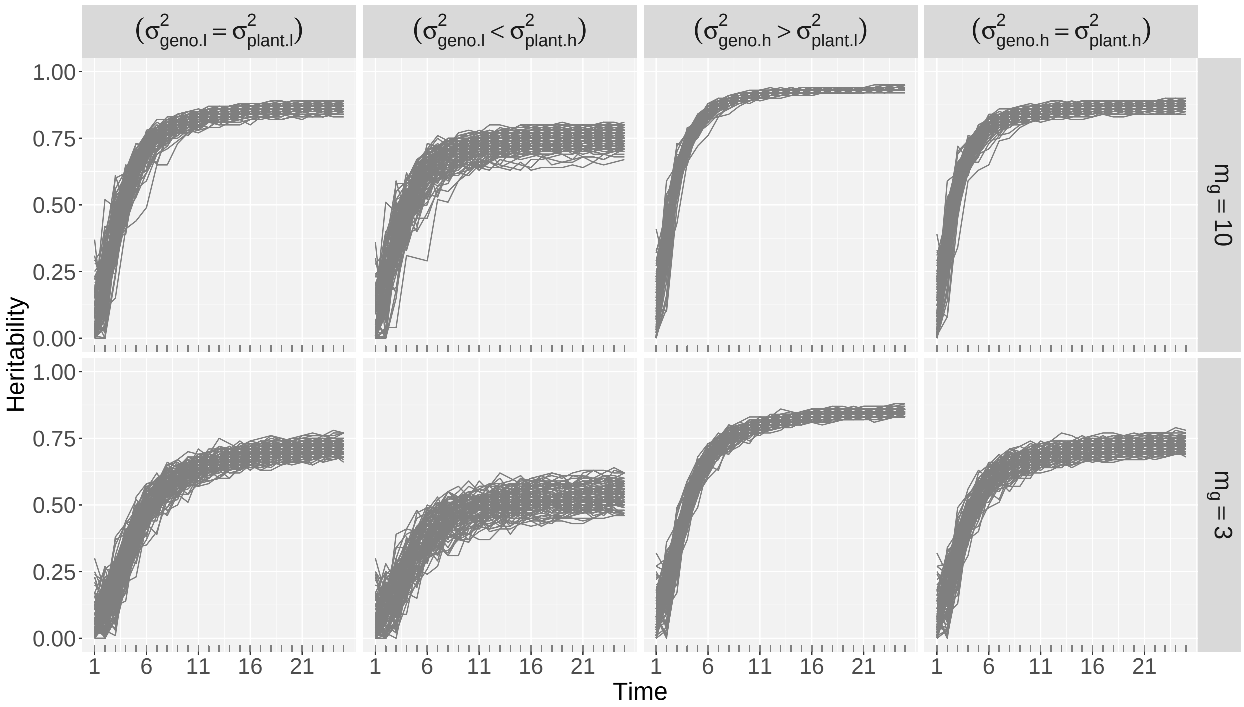


Web Figure 2: Heritability over time for the simulated data under eight simulation scenarios (i.e., $(\sigma^2_{\text{geno}}, \sigma^2_{\text{plant}},\, m_g)$). Each curve corresponds to one simulated dataset (100 datasets by scenario). Heritability is calculated using `SpATS` (Rodríguez-Álvarez et al., 2018) per time point.

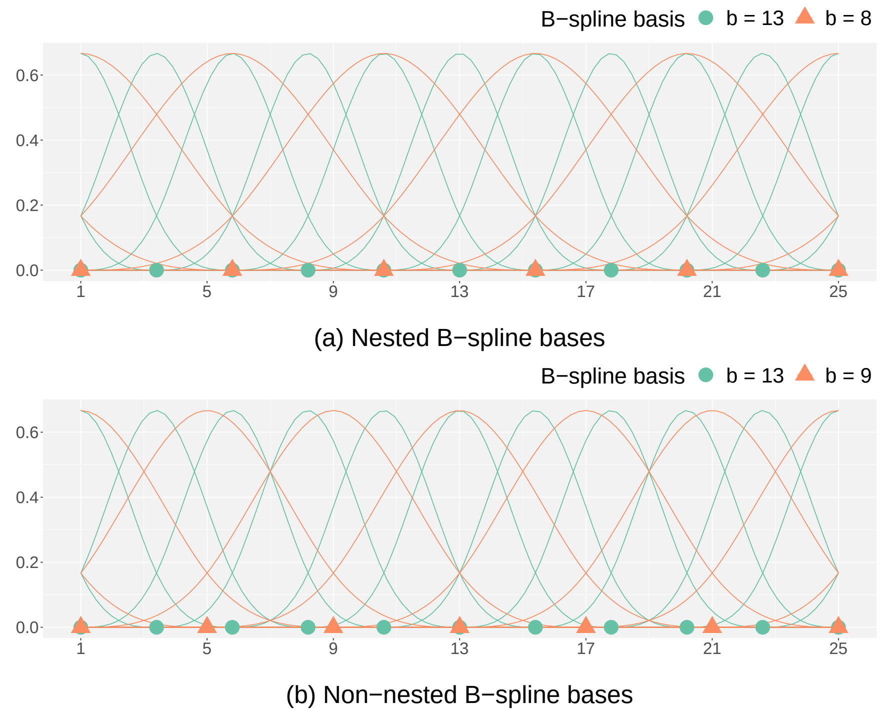


Web Figure 3: Illustrative visualisation of (a) nested (e.g., $(b_1 = 13, b_2 = 8)$) and (b) non-nested B-spline bases (e.g., $(b_1 = 13, b_2 = 9)$).

### Web Appendix C Additional results for the application

Web Figure 7 depicts the same results at genotype level for the PhenoArch platform as shown in Figure 5 of the main text but for all genotypes. In this case, colours represent the one-stage (in blue) and two-stage (in green) approaches.

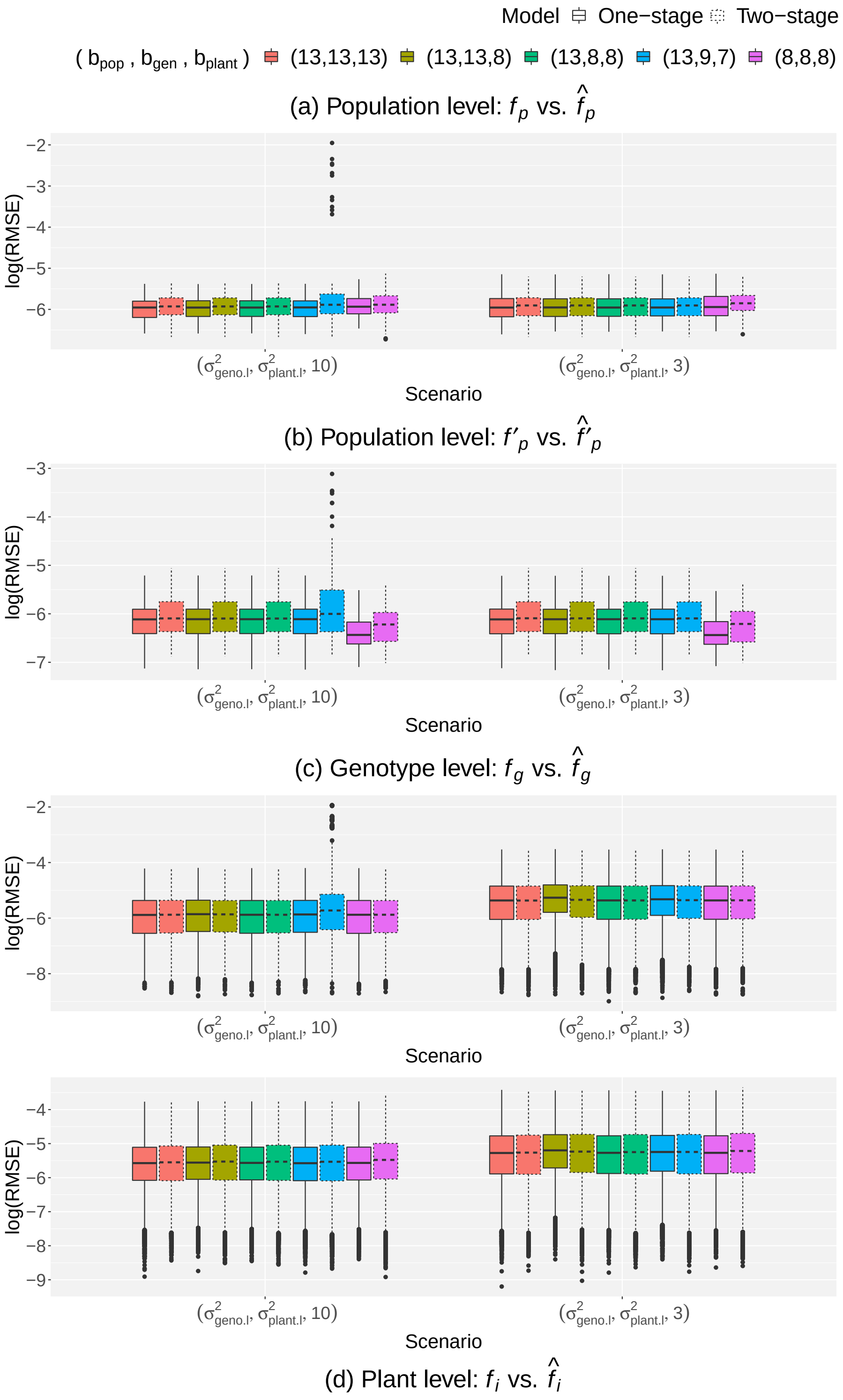


Web Figure 4: Simulation results: Comparison of the simulated and estimated (a) population trajectories, (b) first-order derivative of the population trajectories, (c) genotype deviations curves, and (d) plant deviation curves for two of the eight scenarios of data simulation ($\sigma^2_{\text{geno.l}} = \sigma^2_{\text{plant.l}}$ and $m_g = 3, 10$), using the one- and two-stage approaches, and five B-spline basis configurations ($b_{\text{pop}}$,$b_{\text{gen}}$,$b_{\text{plant}}$) at population, genotype and plant level, respectively.

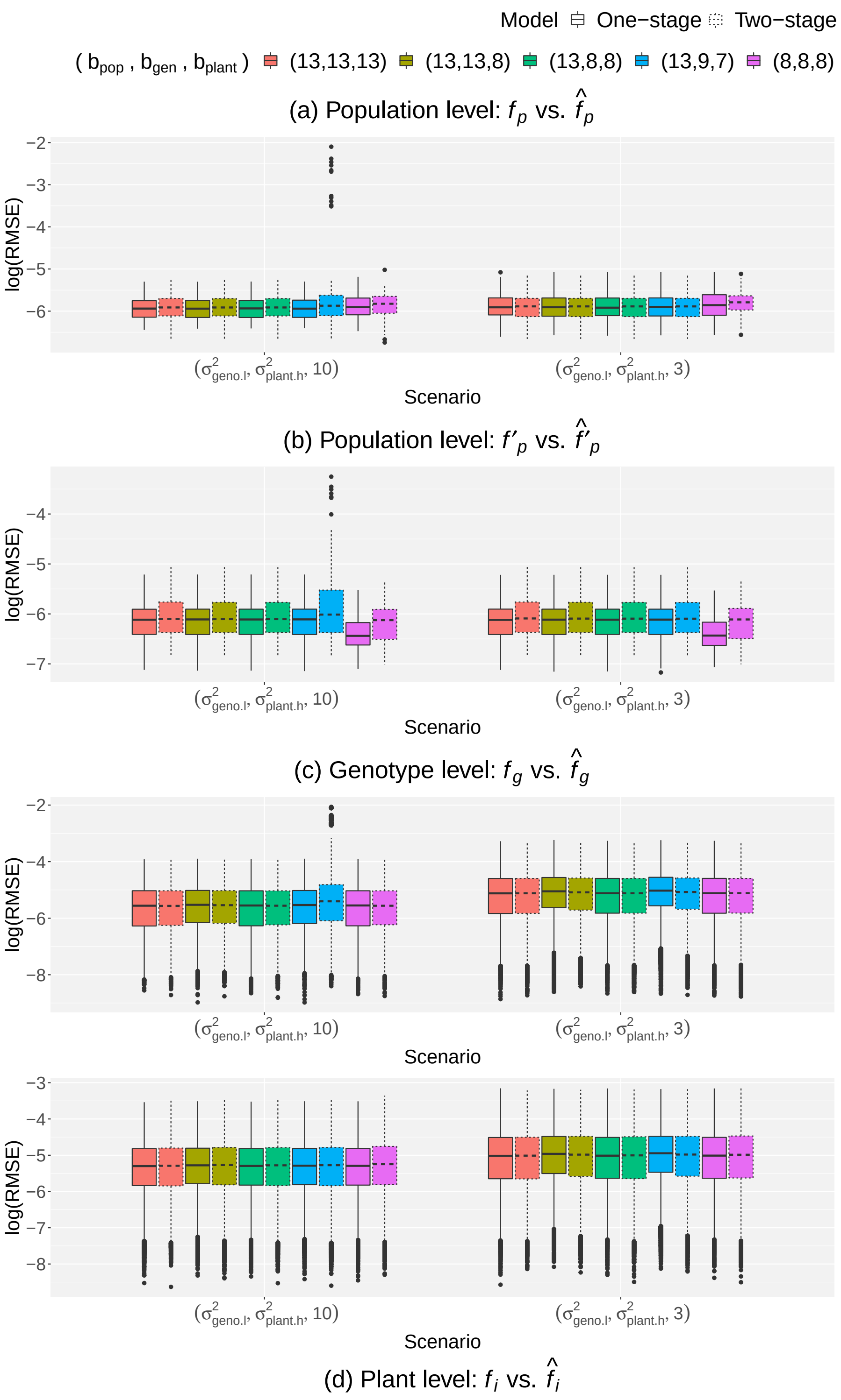


Web Figure 5: Simulation results: Comparison of the simulated and estimated (a) population trajectories, (b) first-order derivative of the population trajectories, (c) genotype deviations curves, and (d) plant deviation curves for two of the eight scenarios of data simulation ($\sigma^2_{\text{geno.l}} < \sigma^2_{\text{plant.h}}$ and $m_g = 3, 10$), using the one- and two-stage approaches, and five B-spline basis configurations ($b_{\text{pop}}$,$b_{\text{gen}}$,$b_{\text{plant}}$) at population, genotype and plant level, respectively.

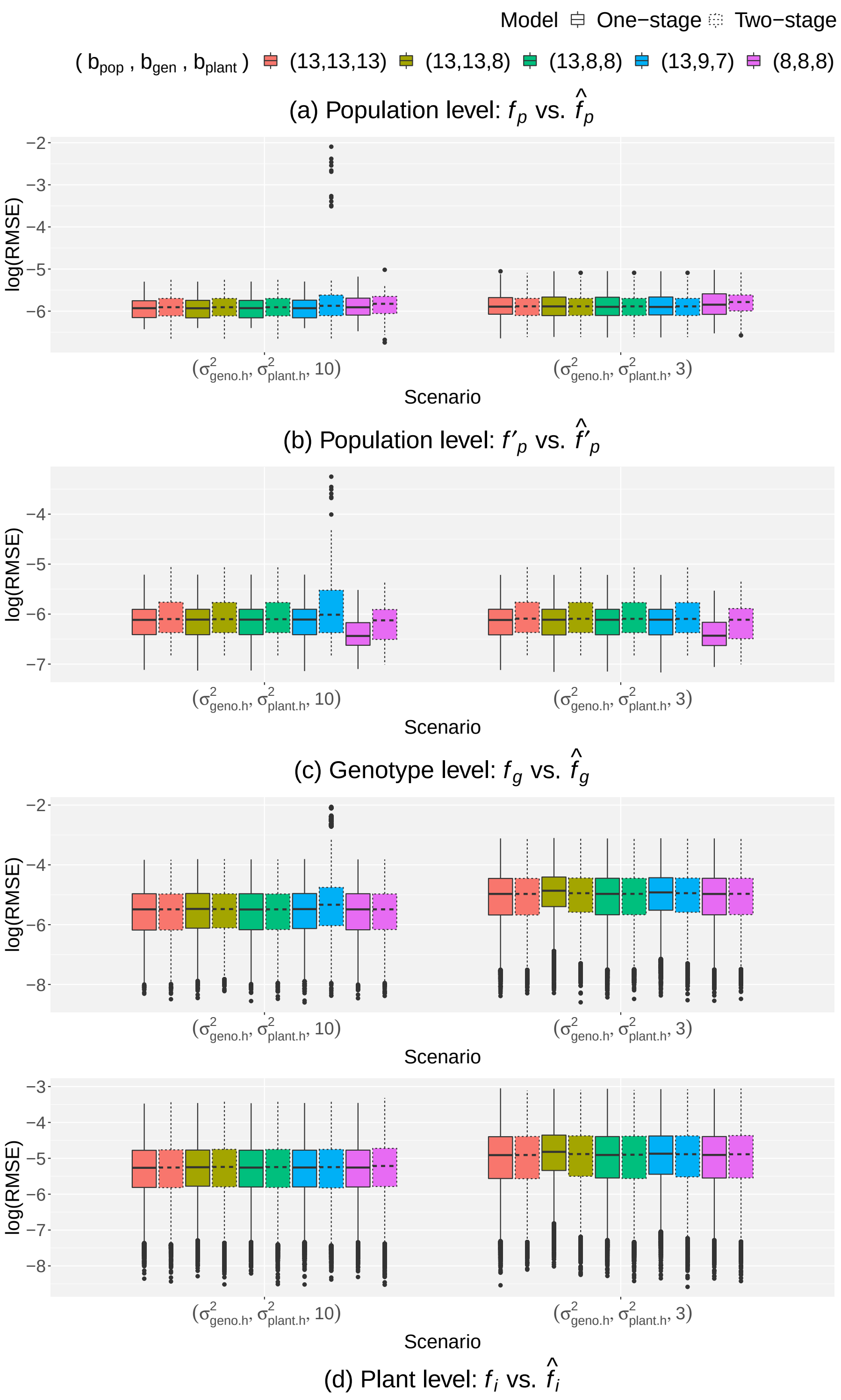


Web Figure 6: Simulation results: Comparison of the simulated and estimated (a) population trajectories, (b) first-order derivative of the population trajectories, (c) genotype deviations curves, and (d) plant deviation curves for two of the eight scenarios of data simulation ($\sigma^2_{\text{geno.h}} = \sigma^2_{\text{plant.h}}$ and $m_g = 3, 10$), using the one- and two-stage approaches, and five B-spline basis configurations ($b_{\text{pop}}$,$b_{\text{gen}}$,$b_{\text{plant}}$) at population, genotype and plant level, respectively.

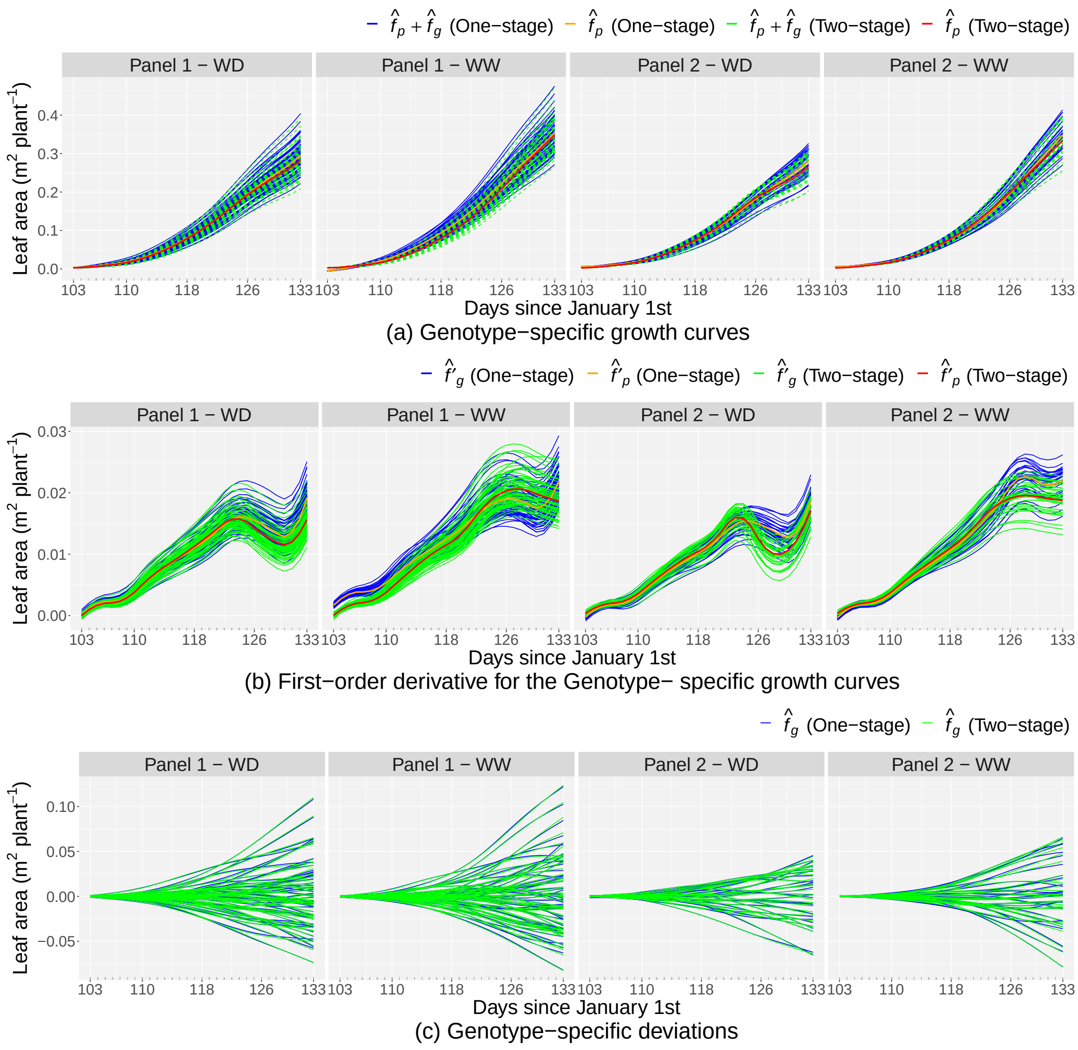


Web Figure 7: Results for the PhenoArch platform at the genotype level (for all genotypes) separately for each population and for both approaches: (a) estimated genotype-specific growth curves, (b) estimated first-order derivative for the genotype-specific growth curves, and (c) estimated genotype-specific deviations. Results for the one-stage approach are in blue, and for the two-stage approach in green. In Figures (a) and (b) orange and red lines represent curves at population level for the one- and two-stage approaches, respectively.
